# Dissecting Hes-centered transcriptional networks in neural stem cell maintenance and tumorigenesis in *Drosophila*

**DOI:** 10.1101/2020.03.25.007187

**Authors:** Srivathsa S. Magadi, Chrysanthi Voutyraki, Gerasimos Anagnostopoulos, Evanthia Zacharioudaki, Ioanna K. Poutakidou, Christina Efraimoglou, Margarita Stapountzi, Vasiliki Theodorou, Christoforos Nikolaou, Christos Delidakis

## Abstract

Neural stem cells divide during embryogenesis and post embryonic development to generate the entire complement of neurons and glia in the nervous system of vertebrates and invertebrates. Studies of the mechanisms controlling the fine balance between neural stem cells and more differentiated progenitors have shown that in every asymmetric cell division progenitors send a Delta-Notch signal back to their sibling stem cells. Here we show that excessive activation of Notch or overexpression of its direct targets of the Hes family causes stem-cell hyperplasias in the *Drosophila* larval central nervous system, which can progress to malignant tumours after allografting to adult hosts. We combined transcriptomic data from these hyperplasias with chromatin occupancy data for Dpn, a Hes transcription factor, to identify genes regulated by Hes factors in this process. We show that the Notch/Hes axis represses a cohort of transcription factor genes. These are excluded from the stem cells and promote early differentiation steps, most likely by preventing the reversion of immature progenitors to a stem-cell fate. Our results suggest that Notch signalling sets up a network of mutually repressing stemness and anti-stemness transcription factors, which include Hes proteins and Zfh1, respectively. This mutual repression ensures robust transition to neuronal and glial differentiation and its perturbation can lead to malignant transformation.

## INTRODUCTION

*Drosophila* neural stem cells (NSCs), a.k.a. neuroblasts, are a well-studied system for stem cell self-renewal, differentiation and carcinogenesis. NSCs are born early in embryogenesis and over the course of embryonic and juvenile (larval, early pupal) life, give birth to all neurons and glia that make up the central nervous system (Lee 2017). They do so by undergoing asymmetric cell divisions that regenerate a NSC and also give rise to a more differentiated cell, which can be either a postmitotic neuron (rare Type 0 mode of division), a ganglion mother cell (GMC, Type I) or an intermediate neural precursor (INP, Type II) (Baumgardt et al. 2014, Homem et al. 2015). The GMC subsequently divides only once to give rise to two postmitotic cells (neurons or glia), whereas the INP can also self-renew, albeit for fewer rounds than a NSC, and generate GMCs. These three modes of asymmetric cell division have also been observed in vertebrate NSCs and the conservation extends beyond cellular phenomenology to the molecular players involved: many proteins that orchestrate NSC divisions and impact on the fate of the progeny cells have mammalian orthologues with similar functions (Paridaen and Huttner 2014). This justifies the use of *Drosophila* to elucidate conserved features of neurogenesis and to pinpoint defects that can lead to runaway proliferation and potential malignant transformation of NSCs, which is the cause for many types of childhood and adult brain tumours.

To ensure the asymmetric outcome of its division, the *Drosophila* NSC polarizes its cortical cytoplasm and aligns the mitotic spindle along this axis (Loyer and Januschke 2019). One of the several consequences of this elaborate mechanism is the unidirectional Delta-Notch signalling from the GMC/INP progeny to the NSC, achieved by asymmetrically segregating Numb, a Notch inhibitor, to the GMC (Knoblich et al. 1995, Spana and Doe 1996). However, Notch signalling is not indispensable for NSC maintenance, as only minor defects are observed upon loss of Notch function: most Type I lineages are unaffected, whereas Type IIs terminate prematurely (Almeida and Bray 2005, Wang et al. 2006, Zacharioudaki et al. 2012). Transcription factors (TFs) of the Hes family are common Notch targets across tissues and species. During *Drosophila* neurogenesis, the NSCs express *dpn* and at least three *E(spl)* genes out of the 11 *Hes* genes in the fly genome (Bier et al. 1992, Almeida and Bray 2005, Zacharioudaki et al. 2012) (see also Figure 1 – supplement 1). Double knockout of *dpn* and the *E(spl) complex* results in acute NSC loss by premature termination of both Type I and Type II lineages (Zacharioudaki et al. 2012, Zhu et al. 2012). Even though *E(spl)* genes as well as *dpn* are induced by Notch, we showed that Dpn expression persists in Notch loss-of-function, justifying the weaker phenotype of the latter. Therefore, the Hes TFs and, to a lesser extent, Notch are central to the maintenance of stemness in fly NSCs. The dependence of NSC maintenance on Hes activity and Notch has also been documented in mammals (Kageyama et al. 2019).

Unlike the mild lof effects of Notch, its overactivation has dramatic consequences on neurogenesis: many immature differentiating cells revert to NSCs, causing extensive NSC hyperplasia at the expense of differentiated neuronal progeny in the larval CNS (Wang et al. 2006, Bowman et al. 2008, Weng et al. 2010, Zacharioudaki et al. 2012). This effect is complete in Type II lineages, but it is also evident in Type Is, albeit with reduced penetrance among lineages. When we overexpressed *dpn* or *E(spl)mγ*, we saw no effects in Type I lineages and only got mild hyperplasias in Type II lineages, much weaker than those caused by Notch overactivation. We later found that Notch also turns on a large number of other targets in the CNS, including temporal TFs, which could contribute by reverting NSC progeny to a more stem-like fate (Zacharioudaki et al. 2012, Zacharioudaki et al. 2016).

Other genetic insults can give rise to NSC hyperplasia in larval CNSs, typified by *brat* or *pros* loss-of-function (Kang and Reichert 2015). Both of these gene products are asymmetrically segregated in the GMC and/or INP (only Brat in the latter, as *pros* is not expressed in Type II NSCs) and function to regulate transcription (Pros) or translation (Brat) (Choksi et al. 2006, Loedige et al. 2015, Reichardt et al. 2018). Compromising the function of the asymmetric cell division machinery (e.g. mutations in *insc, pins, l(2)gl, dlg1*) also often leads to NSC hyperplasia (Gateff 1978, Caussinus and Gonzalez 2005). These hyperplasias are prone to malignant overgrowth and metastasis to distant sites upon allografting to an adult host. It was not known whether the Notch-dependent hyperplasias also have a malignant potential.

Here we revisited the ability of Hes TFs to cause NSC hyperplasia and tested the ability of both Notch and Hes induced hyperplasias to progress to malignant tumours. We show that upon ectopically expressing pairs of Hes transgenes we can elicit severe NSC hyperplasias. These hyperplasias, as well as those caused by Notch overactivation, are malignant upon allografting. Comparison of hyperplastic CNS transcriptomes showed high similarity between two different 2xHes combinations and moderate similarity between these and the high-Notch hyperplasias. Much less similarity was found with other known NSC tumour transcriptomes deposited in databases, pointing to a Hes/Notch-type of tumour. As Hes proteins are known to act as repressors, we focussed on the downregulated targets which are at the same time bound by Dpn by chromatin IP. These were enriched for TFs, among which *pros* and *nerfin1*, which had previously been shown to prevent GMCs and early neurons from reverting to a NSC fate, as well as *erm* which does the same in Type II INPs. We chose to study two other Hes-downregulated TFs, Zfh1 and Gcm, in more detail and will present their expression pattern and possible role in modulating tumorigenesis.

## RESULTS

### Overexpression of Hes genes causes malignant NSC tumours

Overexpression of activated Notch has been known to cause NSC hyperplasia in larvae (Wang et al. 2006) and our earlier work (Zacharioudaki et al. 2012, Zacharioudaki et al. 2016) had shown that this (a) is accompanied by induction of *Hes* genes *dpn* and *E(spl)mγ* and (b) is suppressed by deletion of the *E(spl)* locus. We had also shown that the mild Type II hyperplasias caused by *E(spl)mγ* overexpression bypass the need for Notch activation. To test whether stronger defects can be observed by *Hes* gene overexpression we increased the transgene dose by expressing two copies of Hes genes. We selected to coexpress either *E(spl)mγ + E(spl)m8* or *E(spl)mγ + dpn*, as these are normally expressed in larval NSCs (Figure 1 – supplement 1) and we had earlier shown that these combinations give strong DNA binding as heterodimers in vitro (Koumpanakis 2007, Zacharioudaki et al. 2012). Both global NSC-lineage overexpression by *grhNB-Gal4* and clonal overexpression by *act>STOP>Gal4* were performed. We imaged control and hyperplastic CNSs using NSC markers Dpn, Mira and Wor, NSC/GMC marker Ase, GMC/neuron marker Pros (nuclear), neuron marker Elav and glia marker Repo. Using the *grhNB-Gal4* driver, we observed a large number of ectopic NSCs, not only at their normal superficial location but also invading deeper cortical layers, normally occupied by neurons (Figure 1A-F). At the same time there was a deficit in Elav positive neurons throughout the central brain and thoracic ganglion, regions where *grhNB-Gal4* is expressed. *Hes* overexpression was less disruptive than that of *NΔecd*, an activated form of Notch lacking its extracellular domain: *E(spl)mγ + dpn* overexpressing animals survived to pupation, although they almost never made it to adulthood; in comparison *NΔecd* animals died as early larvae unless *grhNB-Gal4* was kept inactive using a Gal80ts inhibitor until the last day of larval life.

**Figure 1.**
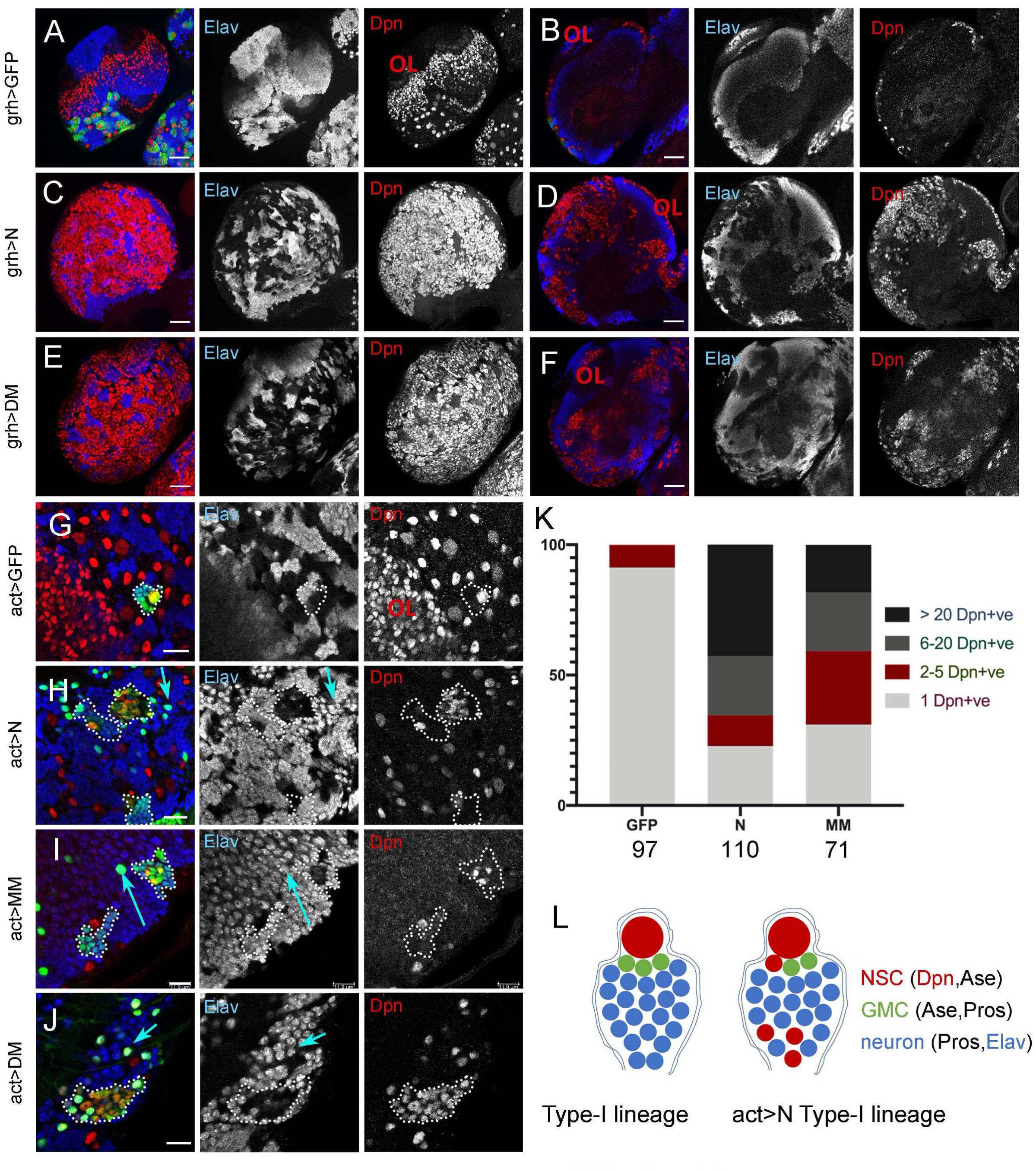
Hes proteins induce neural hyperplasia, similar to Notch. Panels A-J are stained to reveal NSCs (Dpn, red) and neurons (Elav, blue); individual Elav and Dpn channels are also shown in greyscale. (A,B) *grhNB-Gal4; UAS-CD8GFP* brain lobe; (C,D) Brain lobe from *atub-Gal80ts; grhNB-Gal4; UAS-N*Δ*ecd* exposed to the restrictive temperature for 24h; (E,F) Brain lobe from *grhNB-Gal4; UAS-dpn, UAS-mγ*. A,C,E are superficial projections of ∼8 μm; B,D,F are deep single sections at the level of the cortex. (G-J) Single confocal sections of Type-I lineage act>stop>Gal4 FLP-out clones overexpressing GFP (green) (G), GFP + NΔecd (H), GFP + m8 + mγ (I) and GFP + mγ + dpn (J). NSC lineage clones are traced by dotted lines. Single-neuron clones also exist, some examples marked by blue arrows. OL marks the optic lobe, where grhNB-Gal4 is not expressed (A). (K) Quantification of the number of Dpn positive cells per clone in the different genotypes shown in G-I. Numbers below each bar indicate the number of Type I lineage clones scored at 3d after clone induction. The DM condition was excluded from this analysis, since *dpn* is constitutively expressed. (L) Schematics of wt vs NΔecd-overexpressing Type I lineages. DM lineages are similar, with the exception of ubiquitous Dpn expression. MM lineages are also similar, with the exception of frequent detection of lineage cells (GFP positive) that express neither Dpn nor neuronal markers. Scalebars A-F 50μm; G-J 25μm.

Using random clonal expression *(hs-FLP*; *act>STOP>Gal4*) we could focus on the response of individual lineages. We shall henceforth use a short-hand notation for the three effector genotypes to facilitate the discussion: *N* for *UAS-NΔecd, MM* for *UAS-E(spl)mγ+UAS-E(spl)m8* and *DM* for *UAS-dpn+UAS-E(spl)mγ*. In these FLP-out clones, type II lineages were more severely affected (Figure 1 - suplement 2), exhibiting a massive size increase with almost complete absence of differentiating cells (Pros, Elav or Repo positive); only a small number of escapers turned on Pros or Elav; these escapers were more frequent in *DM* and *MM* than in *N* clones. In *N* clones almost all cells expressed the NSC marker Dpn; in *MM* there were many Dpn positive cells, but also many Dpn /Pros doubly negative cells. In *DM* clones all cells were Dpn positive by design (*UAS-dpn* overexpression), although its levels varied greatly from cell to cell, suggesting the existence of post-translational regulation on Dpn protein. Unlike Type II lineages, Type Is showed NSC hyperplasia with variable penetrance and expressivity (Figure 1G-K). About 30% looked wt, whereas 70% contained three or more (up to 176) Dpn positive NSC-like cells; all clones contained differentiating cells (Pros/Elav positive). Phenotype expressivity (number of Dpn positive cells per lineage) was higher in *N* than in *MM* clones. As in Type II, *DM* clones expressed Dpn broadly, albeit at highly variable levels from cell to cell within the same clone. The ectopic NSC-like cells were dispersed and were sometimes located in deep cortical layers, away from the parent NSC, where postmitotic neurons normally reside. Elav and Pros were partially repressed. Elav repression was more effective in *N* than in *MM* or *DM*. As a rule, cells that accumulated high levels of Dpn downregulated Elav, although in *MM* clones Dpn negative/ Elav negative cells were often seen. Nuclear Pros downregulation was less common, in fact many cells were found to co-express Dpn and nuclear Pros in *N* clones (and in *DM* by design, but not in *MM*), suggesting that *N* misexpression turns on Dpn before turning off Pros (Figure 1 – supplement 3). These phenotypes are consistent with a scenario where ectopic Notch or Hes activity causes dedifferentiation of some GMCs and early neurons back to a NSC-like fate. Some of the dedifferentiated clones contain NSCs as late as in pupal or even adult stages (Figure 1 – supplement 4), a time when normal NSCs have ceased proliferating and are not detectable anymore (Maurange et al. 2008, Sousa-Nunes et al. 2010, Homem et al. 2015, Lee 2017). Contrary to the behaviour of NSC lineages, misexpression of *N, MM* or *DM* in post-mitotic mature neurons (located in deep layers, close to the neuropils, or in the abdominal ganglion), did not result in loss of Elav (Figure 1H-J, blue arrows), suggesting that N/Hes activity is effective in causing dedifferentiation only within a certain competence window after neuron birth.

To determine whether the NSC hyperplasias produced by persistent N/Hes activity are malignant we used an allograft assay (Rossi and Gonzalez 2015). In *Drosophila* a tissue is characterized malignant if it can overgrow upon transplantation to an adult host to the point of causing its premature death. We transplanted fragments of CNSs containing *act>STOP>Gal4* FLP-out clones driving *N, MM* or *DM*. Either a brain hemisphere or a ventral nerve cord (VNC) was injected into the abdomen of a young female host. Out of the flies that survived the transplantation process, 60% developed tumours within 20 days (Figure 2). This was seen as increased GFP fluorescence throughout the body, away from the transplantation site. GFP positive tissue dissected out of the body cavity of tumour-bearing flies was retransplanted into new hosts, whereupon tumour growth was observed in almost 100% of injected hosts and this could be repeated for many rounds (data not shown). The high efficiency of tumorigenesis suggests that persistent Hes expression or excessive Notch signalling are sufficient to cause malignant transformation to larval NSC lineages. The fact that we readily recovered transplantable tumours even from VNC allografts suggests that N and Hes can drive malignancies not only from Type II, but also from Type I lineages (data not shown). The tumours recovered from hosts’ abdomens at 10 - 20 days after transplantation expressed nuclear Dpn broadly. Within the tumour mass, several cells were positive for nuclear Pros (GMC/ early neuron-like; Figure 2), whilst Elav positive neuron-like cells were also observed more sparsely (examples shown later in Figure 10). These more differentiated cells had generally low or undetectable levels of Dpn, although double positive cells were also occasionally seen. These results indicate a NSC-like tumour phenotype with little differentiation.

**Figure 2.**
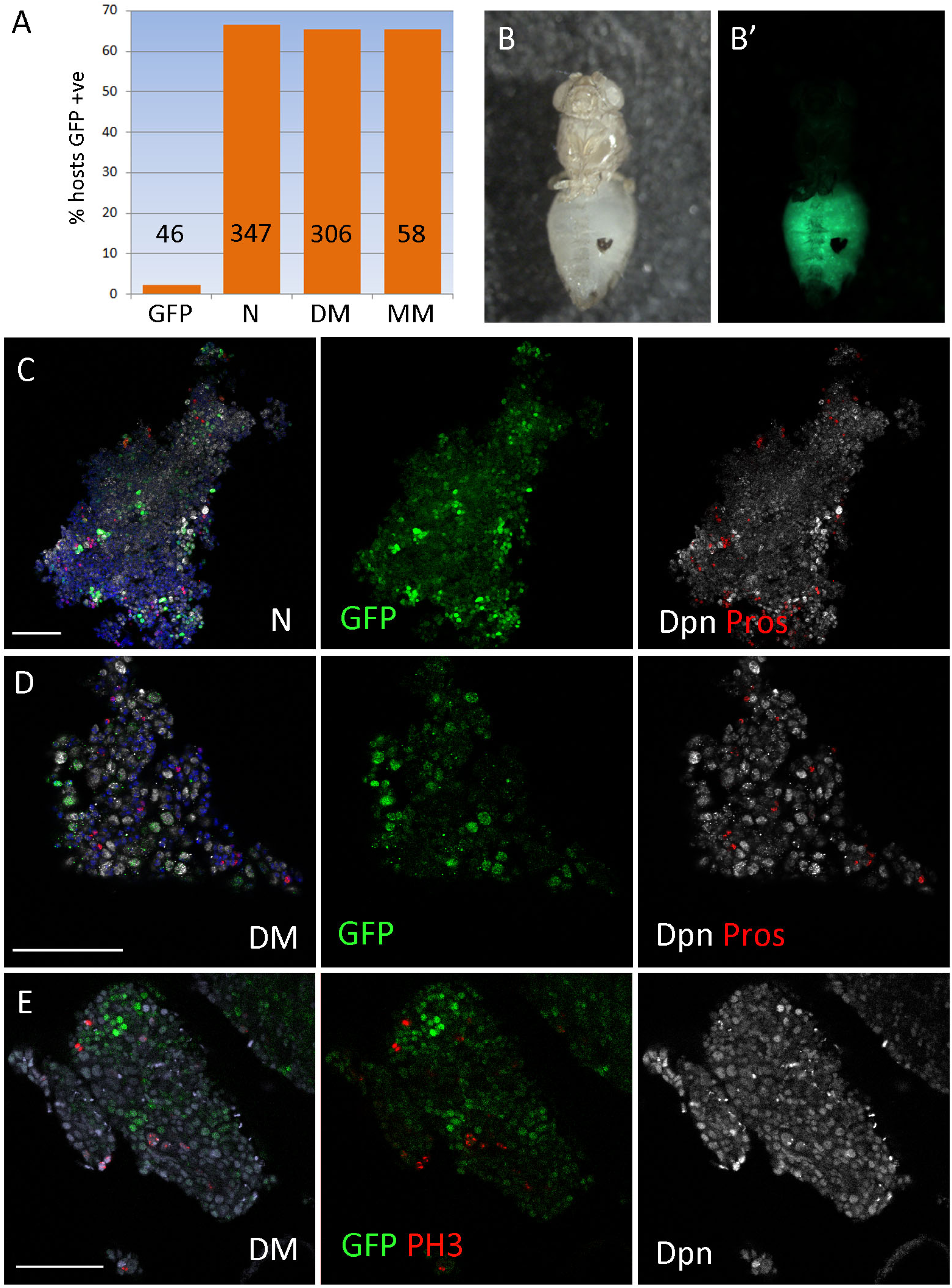
NΔecd and Hes overexpression generates transplantable tumours. Single brain lobes containing *act>stop>Gal4* induced clones overexpressing *N, DM* or *MM* were allografted to adult wild type female abdomens. A shows the percent of hosts that developed GFP positive tumours that expanded in the abdomen within 21 days of transplantation. The numbers in the bars indicate the sample size. One indicative host injected with an act>N brain is shown in B (brightfield) and B’ (epifluorescence). C-E Fragments of tumour recovered from allografted hosts’ abdomens and stained for the markers shown. PH3 refers to the mitotic marker histone H3-phosphoS10. Blue is DAPI. Scale bars 50μm.

### Comparison of transcriptomes from different CNS primary tumours

Since both Hes overexpression and Notch overactivation produce NSC tumours, we wanted to find out to what extent the CNS transcriptomes following these insults are similar or different. For this reason we performed microarray analysis using RNA from hand-dissected CNSs exposed for 24h (by heat-inactivating the the *αtub-Gal80ts* repressor of Gal4) to *DM, MM* or *dpn* (*D*) overexpression via *grhNB-Gal4*. We compared these to N and GFP (control) overexpressing CNSs using the same overexpression regimen (Zacharioudaki et al. 2016). For this analysis (Figure 3) we decided to focus on the N and 2xHes combinations (*DM* and *MM*), since these gave the strongest hyperplasias. Out of 8479 probesets with detectable expression in control larval CNS, we found 1410 that were differentially expressed by at least 1.8x at a FDR ≤0.05 in at least one of these three conditions. *MM* and *DM* showed very similar transcriptomes (Figure 3D) – of the 1410 DEGs, none showed an opposite trend between these two conditions (up in one and down in the other). The N transcriptome was more distinct, but even so, only a very small number of probesets (eight) showed an opposite trend between *N* and *MM/DM* (union of differentially regulated probesets in *MM* and *DM*). Correlation analysis showed that Hes tumours, including the weaker D (*dpn* alone), are more similar to each other than to the Notch tumour (Figure 3D). In conclusion, Hes overexpression causes dramatic transcriptome changes in the CNS, which bears similarities and differences with that caused by NΔecd overexpression. Using RT-qPCR we showed that RNA levels of several genes identified as differentially regulated in the microarrays showed the same trend when individually tested (Figure 3 – supplement 1).

**Figure 3.**
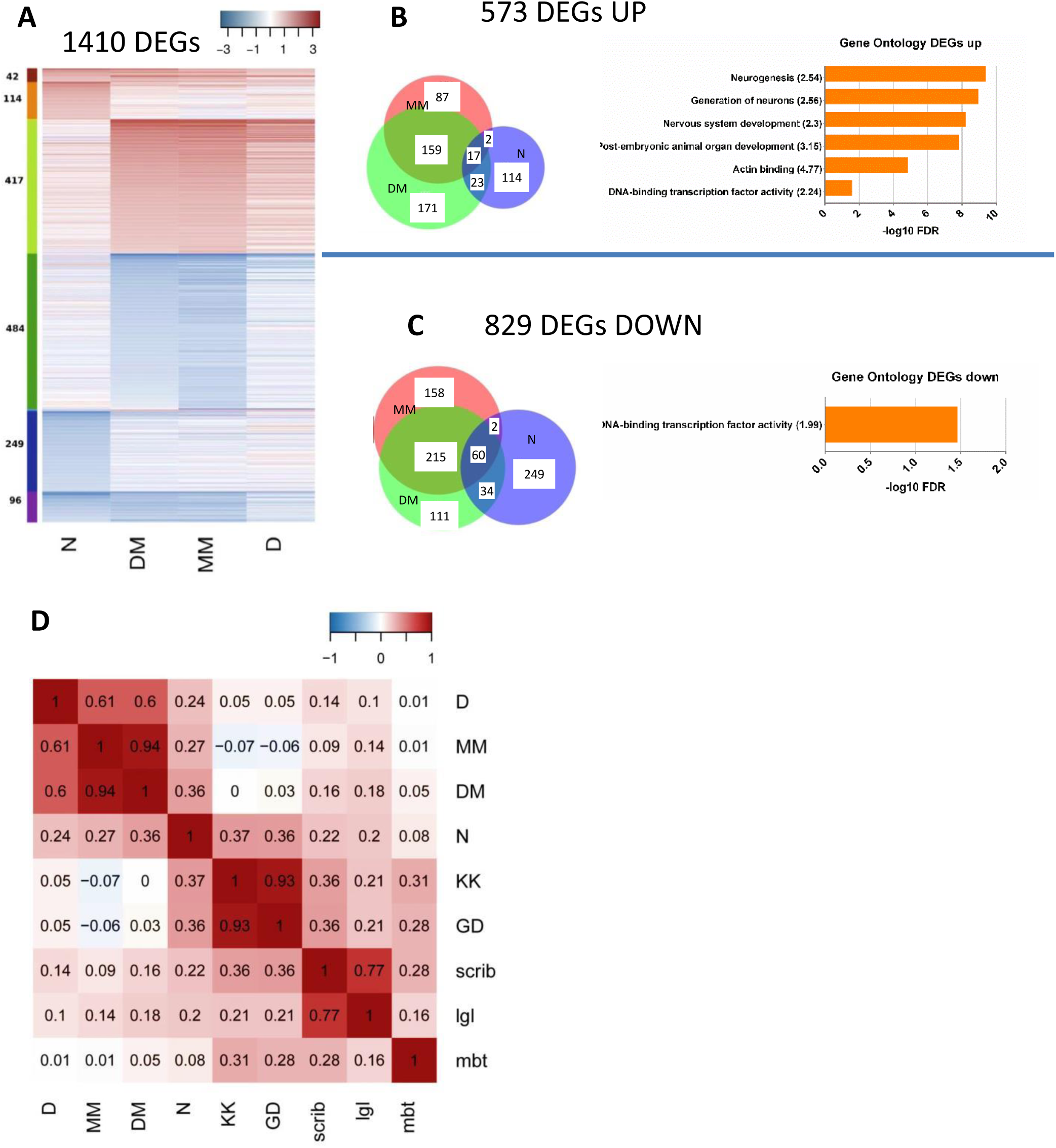
Microarray analysis of N vs Hes overexpressing larval CNSs. **(A)**. Heatmap showing the log_2_(fold-change) of all 1410 genes (probesets) whose expression changed by at least 1.8x compared to wild-type CNS controls (FDR ≤ 0.05) in at least one of the four conditions (N, DM, MM or D expressed by grhNB-Gal4 for 24h). (**B**,**C)**. The 1410 differentially expressed genes (DEGs) were split into 573 upregulated (B) and 829 downregulated (C) (and 8 oppositely regulated; not shown). The Venn diagrams classify these genes according to the condition(s) where they showed change with high confidence (FDR ≤ 0.05). Note that the overlaps would have been much broader, if we had not imposed this strict FDR-value cutoff. GO terms showing significant enrichment in these gene sublists are also shown with fold enrichment in parentheses. **(D)** Pearson correlation coefficients of the distributions of fold-change values for nine CNS tumour transcriptomes, calculated on the basis of the entire set of 8479 genes showing detectable expression in our control arrays, whether they were differentially regulated or not. scrib, lgl and mbt refer to homozygous mutant CNSs for *scrib, l(2)gl* and *l(3)mbt*; KK and GD refer to two *brat* RNAi lines driven by *insc-Gal4*.

To address whether our transcriptome changes simply reflect the change in cellular composition of the CNSs used, we asked whether they correspond to upregulation of NSC specific genes and/or downregulation of neuron specific genes. When we queried a list of genes differentially expressed in wt NSCs vs neurons (Berger et al. 2012), no such trend was observed. Instead, most neuron- and NSC-enriched genes were unchanged in the N/Hes CNSs and conversely most N/Hes differentially regulated genes belonged to neither the neuron nor the NSC-enriched gene-sets (Figure 3 – supplement 2). The fraction of N/Hes differentially regulated genes that overlapped the NSC/neuron gene-sets displayed changes in both directions (up and down) regardless of neuron or NSC enrichment. Similar results were obtained with a more recent categorization of genes expressed differentially between (wt) NSCs and GMCs (Wissel et al. 2018). We conclude that the transcriptomic changes that we documented are mostly due to the deregulation of the Notch/Hes axis and less due to the accompanying over-representation of NSCs vs GMCs and neurons.

Other genetic insults have been reported to generate brain tumours in *Drosophila* (reviewed in (Kang and Reichert 2015)). We recovered transcriptome data for two tumours formed by *brat* RNAi in NSC lineages (using *insc-Gal4*) (Neumuller et al. 2011), two caused by the loss of cell polarity genes *l(2)gl* and *scrib* (GEO accession GSE48852) and one caused by the loss of the nuclear architecture gene *l(3)mbt* (Janic et al. 2010). When we compared these with our N/Hes induced tumours (Figure 3D, Figure 3 – supplement 3), we saw little or no similarity. Essentially no correlation was observed between the Hes transcriptomes and any of the *brat* RNAi, *l(2)gl*^*-/-*^, *scrib*^*-/-*^ or *l(3)mbt*^*-/-*^. In contrast, the N transcriptome showed a moderate correlation with the two *brat* RNAi conditions, but not the *l(2)gl*^*-/-*^, *scrib*^*-/-*^ or *l(3)mbt*^*-/-*^. This is consistent with the fact that the N/Hes tumours consist mostly of NSC-like cells, whereas the *l(2)gl*^*-/-*^and *scrib*^*-/-*^ tumours strongly express differentiation markers (Beaucher et al. 2007). *l(3)mbt* tumours stood apart from all others, in agreement with a different cellular origin (optic lobe precursors, (Richter et al. 2011)). In short, N overactivation partially resembles Hes overactivation and partially resembles *brat* lof, which are distinct from each other.

Upon GO term analysis of the N/Hes transcriptomes, few terms showed statistically significant enrichment (Figure 3B, C), mostly associated with neurogenesis. Interestingly, “DNA binding transcription factor”, was enriched both in the upregulated and the downregulated sub-list of differentially expressed genes, which prompted us to focus on transcription factor genes for the followup analysis.

### Genome-wide binding of Dpn

Transcriptomic analysis of complex tissues is inherently noisy from the contribution of numerous cell types. In order to reveal the direct targets of Hes proteins in the NSCs we performed chromatin IP using an anti-Dpn antibody on hyperplastic larval CNSs overexpressing *UAS-dpn + UAS-E(spl)mγ* driven by *grhNB-Gal4*. This enabled us to focus on the expanded population of NSCs, since there is little Dpn expression (endogenous or transgenic) in other cell types. From deep sequencing of immunoprecipitated chromatin from two biological replicates we obtained a high confidence set of 229 peaks displaying Dpn occupancy (Figure 4A). Mining these 229 ChIP peaks for over-represented DNA motifs, we obtained an E_B_/E_C_ box sequence and variants as the top hits (Figure 4B), confirming that in an in vivo context Dpn binds sequences that conform to the well described in vitro binding consensus for Hes proteins (Jennings et al. 1999, Winston et al. 1999).

**Figure 4.**
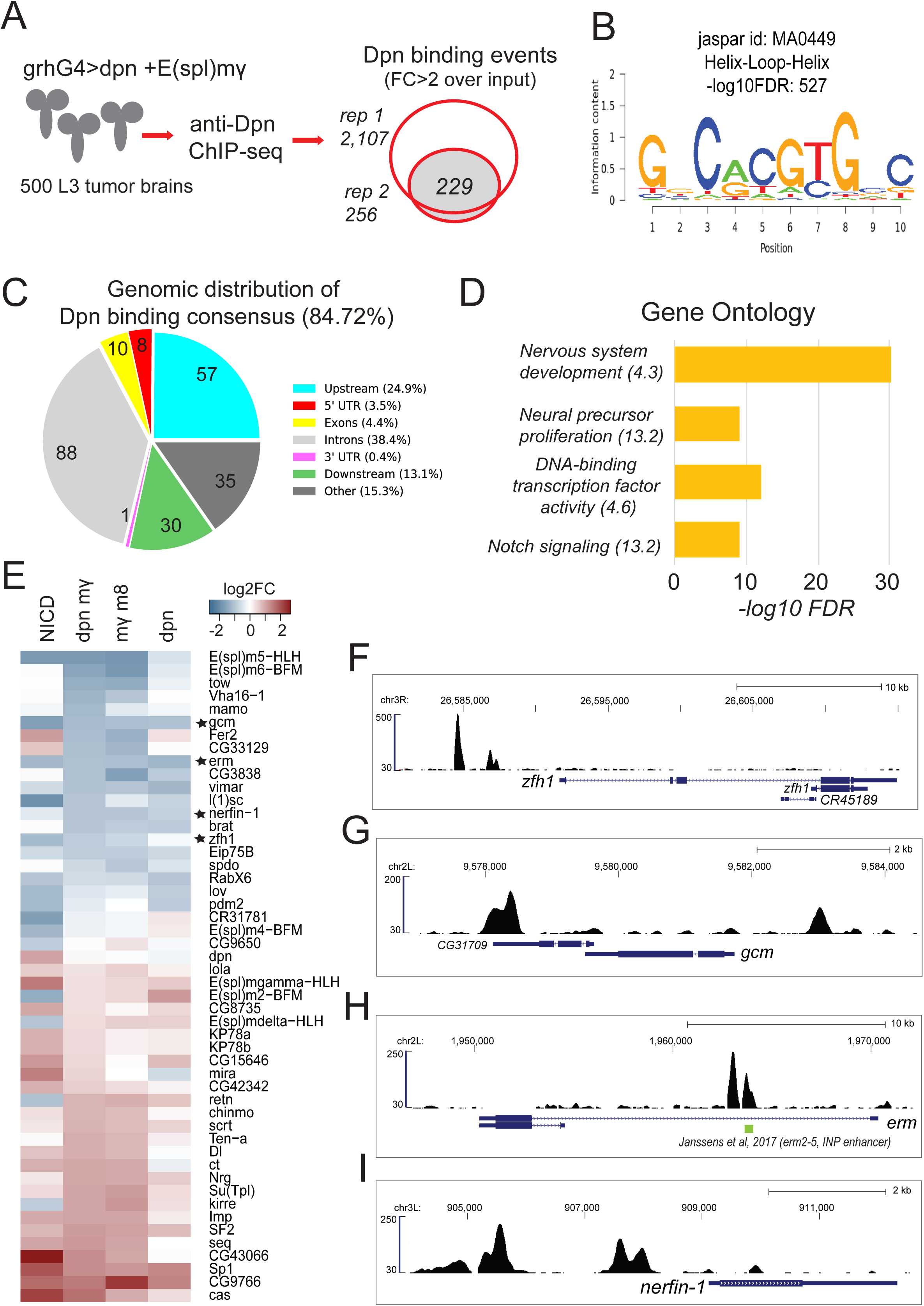
Genome-wide mapping of Dpn binding in *DM* CNS hyperplasias. (A) Chromatin extracts from 500 L3 DM CNSs were isolated and incubated with anti-Dpn bound beads. Two biological ChIP-seq replicates were performed. A Dpn binding consensus of 229 binding events was generated from regions enriched at least 2x input in both replicates. (B) Motif analysis of the Dpn bound consensus revealed a high enrichment for a Hes-type bHLH binding motif. (C) Genomic distribution of Dpn bound consensus peaks. (D) Gene ontology enrichment analysis of the 358 genes with Dpn consensus binding events in their vicinity. (E) Transcriptional response of 50 Dpn-bound genes that were also differentially regulated in the microarray analysis of Figure 3A. (F) Genomic snapshot of Dpn binding at the *zfh1* locus. (G) Genomic snapshot of Dpn binding at the *gcm* locus. (H) Genomic snapshot of Dpn binding at the *erm* locus. (I) Genomic snapshot of Dpn binding at the *nerfin-1* locus.

A large percentage of the Dpn peaks fell within or near genes (Figure 4C). We extracted a list of 358 genes that contain at least one Dpn binding event within a 5kb window up- and down-stream. These genes are enriched for factors involved in neural precursor development, Notch signalling and transcriptional regulation (Figure 4D). When this list of Dpn binding-associated genes was overlapped with the differentially expressed genes from the microarray experiment, we obtained 50 genes, whose transcriptional response is shown in Figure 4E. Although Hes proteins are thought to act as repressors, only half of these 50 Dpn-bound genes showed repression in the Hes-induced tumours. This is not surprising, because of the heterogeneity of the tissue analysed, which means that genes bound by Dpn in the NSCs may also be expressed in other cell types, so their behaviour in the microarray will reflect the average across all cell types in the tissue. Another reason is that mRNA levels respond to multiple transcriptional/ post-transcriptional regulatory inputs, so that the overall effect of Hes overexpression (integrating direct and indirect effects) may not be repression, despite the increased binding of Hes repressors near these genes. We do not exclude the possibility that Hes proteins might behave as activators in certain enhancer contexts, although we have no evidence in support of such a scenario. We focussed our subsequent experimental validation on a set of transcription factor genes which appear to be prominent Hes targets, since they combine high Dpn occupancy with transcriptional repression in the Hes-induced tumours.

### Differentiation-promoting transcription factors are among the most highly downregulated Hes target genes

Amongst the top downregulated transcription factors we noted two zinc-finger genes *erm* and *nerfin1*, known to promote neural differentiation (Figure 4E). Both of these genes have nearby Dpn ChIP peaks implying direct repression by Hes proteins (Figure 4H, I). *erm* is turned on in the immature INP progeny of Type II NSCs and restricts their self-renewal. Its loss of function is sufficient to cause dedifferentiation of INPs to highly proliferative ectopic Type II NSCs (Weng et al. 2010, Janssens et al. 2014, Koe et al. 2014). An enhancer recapitulating its expression was shown to bind stem cell repressors, including Dpn (Janssens et al. 2017) and coincides with one of the two peaks we detected (Figure 4H). *nerfin1* is expressed in newly born neurons in both Type I and II lineages as well as in terminating NSCs after pupariation. Its loss can cause dedifferentiation and overproliferation (Froldi et al. 2015).

Besides, the well-studied *erm* and *nerfin1*, there were another 13 TFs among the 27 downregulated and Dpn-bound TF genes. We turned our attention to two of these, *zfh1* and *gcm* (Figure 4F,G), to ask whether their repression by the N/Hes system might play a role in NSC tumorigenesis. *zfh1* encodes a large protein with a homeodomain and 9 zinc-fingers; it has a broad tissue distribution and pleiotropic functions. In the embryo it participates in the differentiation of various mesoderm derivatives as well as of certain types of CNS neurons and glia, most notably motor neurons (Postigo et al. 1999, Layden et al. 2006, Sellin et al. 2009). In third instar larval and early pupal CNSs we detected moderate expression in newly-born neurons in most brain (Type I and II) and VNC lineages (Figure 5) using an antibody and a *lacZ* enhancer trap (Lai et al. 1991, Justice et al. 1995). Stronger expression was detected in dispersed older neurons located in deeper cortical layers (Figure 5 – supplement 1). Additionally, a large number of glia, perhaps all glia, exhibited detectable *zfh1* expression. Hyperplastic brains generated by overexpression of *NΔecd* or *2xHes* during late larval stages using *grhNB-Gal4* showed a severe reduction of *zfh1* expression in the superficial layers, and a more moderate reduction of *zfh1* positive neurons in deeper layers (Figure 5). The disorganization of the brain and shrinking of the neuropils did not allow us to assess the effect on glial cells.

**Figure 5.**
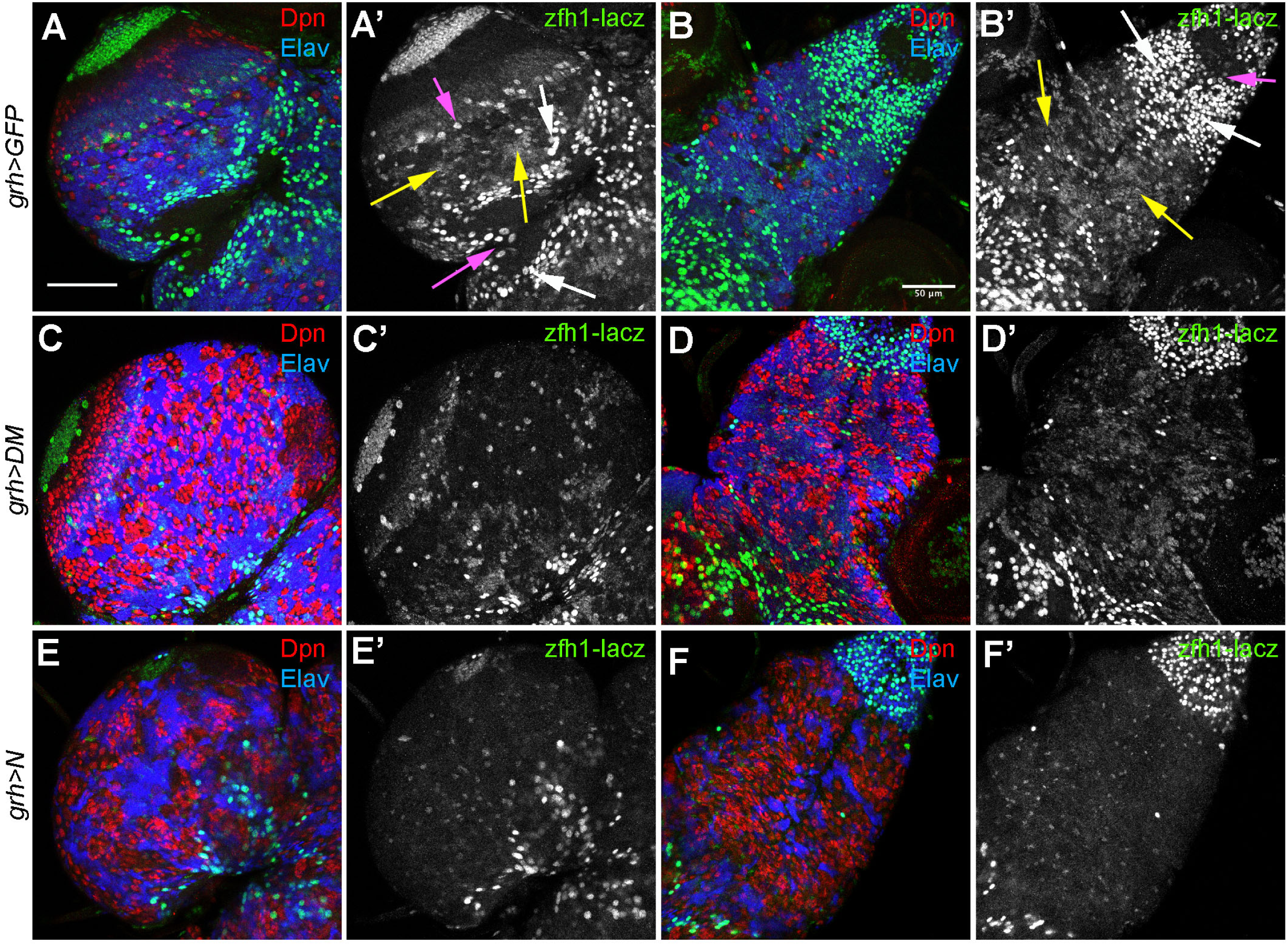
*zfh1* expression in wt and hyperplastic CNSs. *zfh1-lacZ*^*00865*^ was used as a reporter for *zfh1* expression. β-galactosidase is green, Dpn (NSCs) is red and Elav (neurons) blue. A, C, E show brain lobes; B, D, F show VNCs. All CNSs carry *grhNB-Gal4* and (A,B) *UAS-GFP* (wt, GFP not imaged), (C,D) *UAS-dpn, UAS-E(spl)mγ* (DM) and (E,F) *UAS-NΔecd* (N). In the wt panels examples of *zfh1*-expressing GMCs/young neurons are marked with yellow arrows, mature neurons with white arrows and glia with magenta arrows. Scalebars 50μm.

*gcm* encodes a transcriptional activator and founder of the Gcm-family of proteins, which are known to promote gliogenesis (Ragone et al. 2003). *gcm* is transiently expressed in glia precursors and eventually shut off in mature glia; it is also involved in the differentiation of some lamina neurons, embryonic tendon cells and haemocytes (Cattenoz et al. 2016). In the larval CNS we detected *gcm* expression using the *gcm*^*rA87*^*-lacZ* enhancer trap (Jones et al. 1995) (Figure 6). Besides widespread expression in the optic lobe, there was sparse expression in the larval CNS. In deep areas (near the neuropils) LacZ was detected in some Repo positive glial cells as well as nearby cells that could be neuropil glia precursors. At the superficial levels of the central brain and thoracic ganglion, we did not detect LacZ expression in NSCs, but we did detect it in several immature neural precursors (INPs) in both lateral and medial Type II NSC lineages (data not shown). This is in agreement with the fact that some of the Type II lineages produce glia as well as neurons (Viktorin et al. 2011, Viktorin et al. 2013). *gcm-lacZ* was also prominently expressed in two neuronal clusters, one on the ventral and one on the dorsal side of each brain hemisphere – these are Type I derived neurons, but nearby NSCs were *gcm-lacZ* negative (Soustelle and Giangrande 2007). The only NSCs expressing *gcm-lacZ* were found at the medial edge of the medulla and at the optic lobe’s inner proliferation centre (data not shown), probably gliogenic NSCs (Li et al. 2013). Upon *grhNB-Gal4* overexpression of *2xHes* or *<u>NΔecd</u>*, central brain expression of *gcm-lacZ* was downregulated in both neuron and glial precursors (Figure 6), with the exception of some deep glial precursors that retained *gcm-lacZ* expression. Optic lobe expression was unaffected, since *grhNB-Gal4* does not drive expression in the optic lobe.

**Figure 6.**
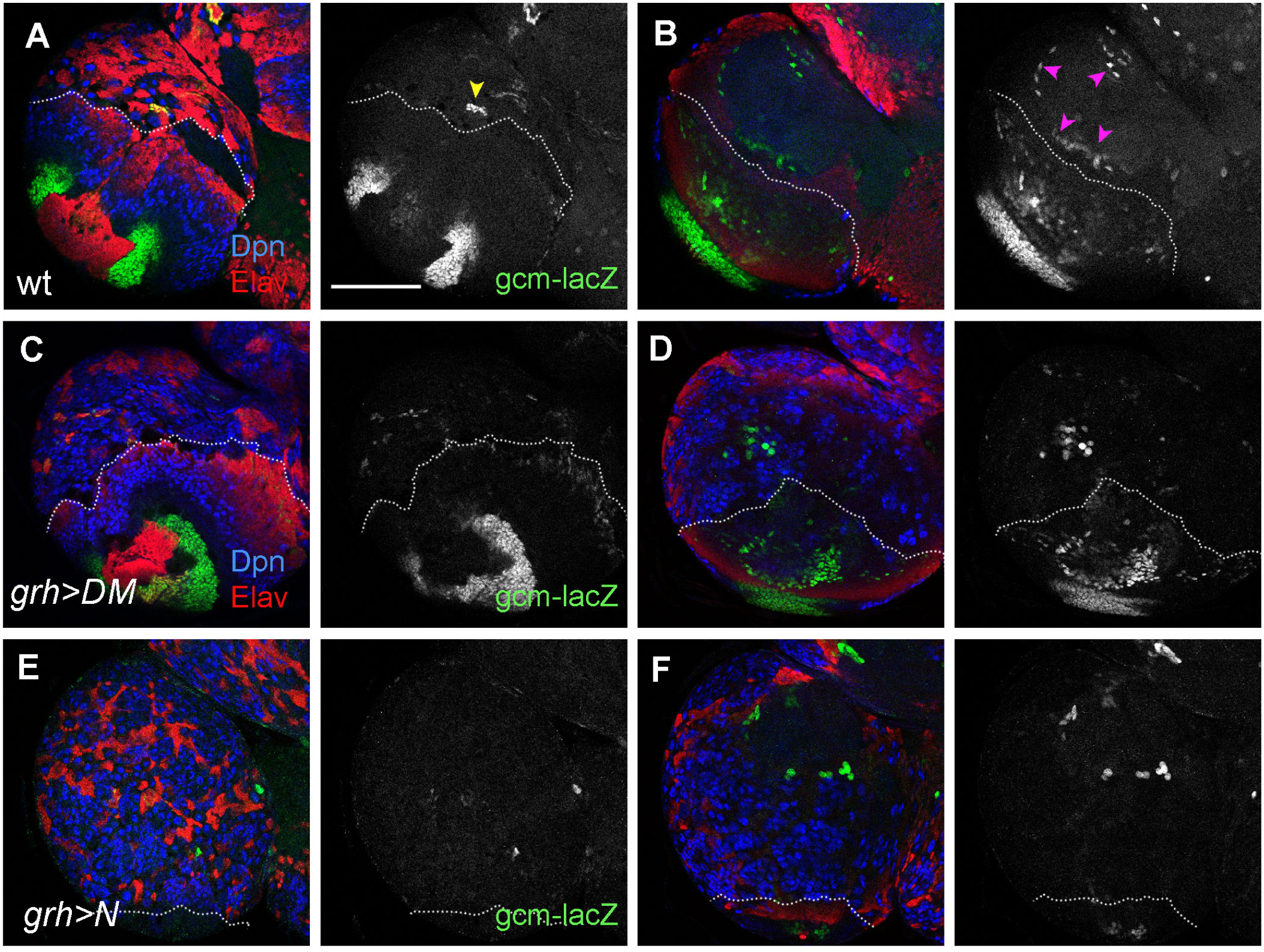
*gcm* expression in wt and hyperplastic CNSs. *gcm-lacZ*^*rA87*^ was used as a reporter for *gcm* expression. β-galactosidase is green, Dpn (NSCs) is blue and Elav (neurons) red. (A,B) A wt brain lobe, superficial (A) and deep (B) section. Glial cells lining the neuropil are indicated by magenta arrowheads. One of the Gcm-positive neuronal clusters is marked by a yellow arrowhead. (C,D) A DM-overexpressing brain lobe, superficial (C) and deep (D) section. (E,F) A N-overexpressing brain lobe, superficial (E) and deep (F) section. In all panels the dotted line indicates the border between central brain (upper) and optic lobe (lower). Scalebar 100μm.

### Loss- and gain-of-function analysis of novel Hes targets

Both embryonic and post-embryonic phenotypes of the loss of *gcm* and *zfh1* have been described. At the larval stage *gcm* lof is known to result in the absence of most glial cells born post-embryonically (Awasaki et al. 2008). In the postembryonic CNS, *zfh1* lof has been studied in the context of specific leg motorneuron lineages and it results in defects in the morphology of some neurons, but not defects in their birth or neurotransmitter phenotype (Enriquez et al. 2015). To address their need in NSCs and their immediate progeny, we generated MARCM clones for null alleles (*zfh1*^*75*.*26*^ or *gcm*^*N7-4*^) 24-48h ALH and studied their effects 3-5 days later, namely in late larval or early pupal (12-24h APF) stages. Unlike *erm* and *nerfin1*, whose loss causes supernumerary Type II and Type I NSCs, respectively, knocking out either *zfh1* or *gcm* had no detectable effect in the cellular composition of either Type I or Type II lineages based on NSC, GMC and neuron markers (Dpn, Pros, Elav; Figure 6 – supplement 1).

We then turned to a gof approach to ask what would happen if we were to forcibly express these Hes-repressed factors in NSCs. For this reason we used an *act>STOP>Gal4* driver and induced expression by *hs-FLP* in mid-larval stages. *zfh1* overexpression in type I lineages resulted in complete elimination of Dpn within 1 day of clone induction (Figure 7). This was accompanied by shrinkage of the NSC, making it in most cases (76% of the clones) indistinguishable from other cells in the clone. Ase, another NSC (and GMC) marker, was not similarly repressed within the first day, although it too was extinguished at later times after clone induction (ACI) (Figure 7 – supplement 1). Interestingly, Elav and Pros were also partially repressed in *zfh1* overexpressing lineages: several cells lacked both Dpn and these differentiation markers in a large proportion of clones (Figure 7H). For comparison, wt clones never contained doubly Dpn/Pros negative cells; nuclear Pros was detectable in all NSC progeny (GMCs and neurons) and Elav gradually came on in neurons, even those that are still close to the NSC at the CNS surface. Thus overexpression of *zfh1* was able to antagonize both NSC and neuron cell fates. These cells were not converted to glia, as the glial marker Repo was not induced in *act>zfh1* clones (Figure 7 – supplement 1).

**Figure 7.**
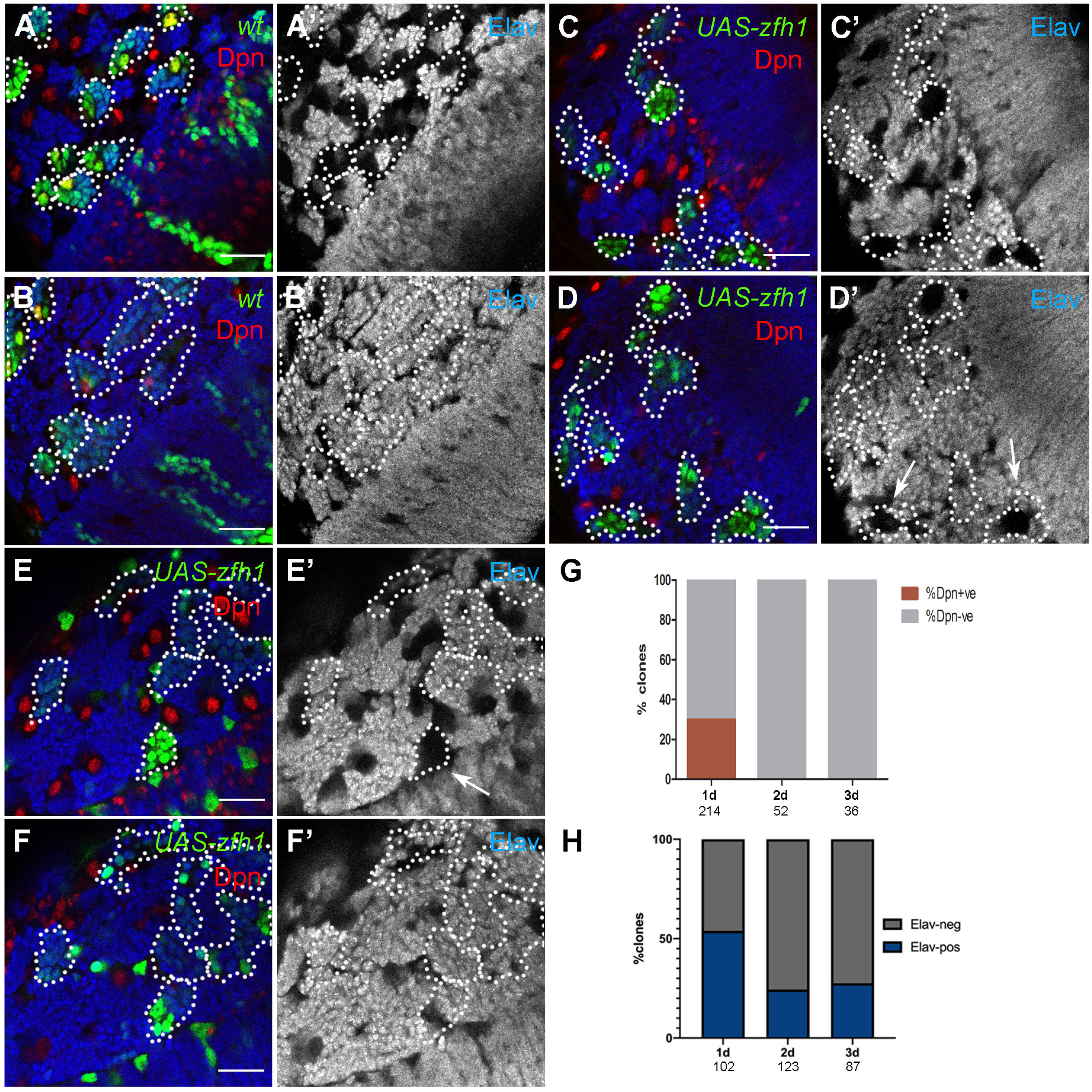
Effects of misexpressing *zfh1* on CNS lineages. *act>STOP>Gal4* FLPout clones are marked by GFP (green) and traced with dotted outlines. NSCs are marked by Dpn (red – yellow when overlapped with green) and neurons by Elav (blue). The Elav channels are shown separately in greyscale. (A,B) Control (wt) clones, 2d after clone induction (ACI) – the same brain lobe area imaged superficially, at the level of NSCs (A) and deeper (B). (C,D) UAS-zfh1 clones 1d ACI; the same brain lobe area imaged superficially (C) and deeper (D). (E,F) UAS-zfh1 clones 2d ACI; the same brain lobe area imaged superficially (E) and deeper (F). Scalebars 25μm. (G) The frequency of UAS-zfh1 clones containing a Dpn positive cell (brown) or lacking Dpn staining entirely (grey) as a function of time elapsed ACI. (H) The frequency of clones showing extensive Elav repression (grey) or exhibiting normal Elav in all cells but the NSC/GMCs (blue) as a function of time elapsed ACI. Arrows in panels D’ and E’ point to examples of Elav-repressed clones.

Unlike *zfh1, gcm* overexpression did not abolish the NSCs. Each marked lineage contained a single Dpn positive NSC, as in control (GFP-only expressing) clones. Also Elav and Pros were not visibly affected. However, a few Repo positive cells were observed in approximately 50% of the overexpressing lineages (Figure 7 – supplement 2), indicative of ectopic gliogenesis, as described by Viktorin et al (2011). Therefore, ectopic expression of *gcm* for 2-3 days does not greatly upset the execution of NSC lineages, other than converting some of the differentiated progeny into glia, consistent with its well-known glia-promoting role.

### Overexpression of Hes targets ameliorates the tumour phenotype

We next wanted to address whether overexpression of *zfh1* or *gcm* would influence Notch/Hes-induced tumorigenesis. If the depletion of these Hes targets contributes towards tumorigenesis in Hes/Notch CNS tumours, perhaps expressing them transgenically would reverse this effect. We therefore co-expressed either *gcm* or *zfh1* with tumour-promoting transgene combinations, namely *N, MM* or *DM*, in random *act>STOP>Gal4* clones.

*zfh1* overexpression in a tumour background produced round clones with a conspicuous absence of Dpn in most lineages (unless *UAS-dpn* was used); only a few scattered escaper Dpn positive cells were seen (Figure 8). Therefore, the ability of Zfh1 to repress *dpn* overrides the latter’s activation by N or MM. Another NSC marker, Ase, responded more variably: it was fully repressed in *UAS-N+zfh1* clones, whereas it showed variable penetrance in *UAS-DM* (or *MM*) + *zfh1* clones (Figure 8 – supplement 1): in some clones it was repressed, whereas in others it was activated in many cells. More remarkable was the strong repression of Elav (Figure 8) and Pros (Figure 8 – supplement 1) in *UAS-N* (*DM, MM*) +*zfh1* clones, which was seen with full penetrance (in all clones) in the majority of the marked cells. Whereas both *UAS-N* (or *DM* or *MM*) and *UAS-zfh1* produced sporadic loss of these differentiation markers (Figures 1 & 7), their combination was far more potent. We excluded the possibility that the *N/Hes+zfh1* expressing cells were massively converted to glia using the pan-glial Repo marker (Figure 8 – supplement 1). In summary, zfh1 overexpression overrides Dpn induction by N/Hes, but synergizes with the latter to maintain CNS progenitors in an undifferentiated state (which sometimes retain the Ase marker), generating cell clumps that stay separate and do not integrate with the cortex of differentiated neurons/glia.

**Figure 8.**
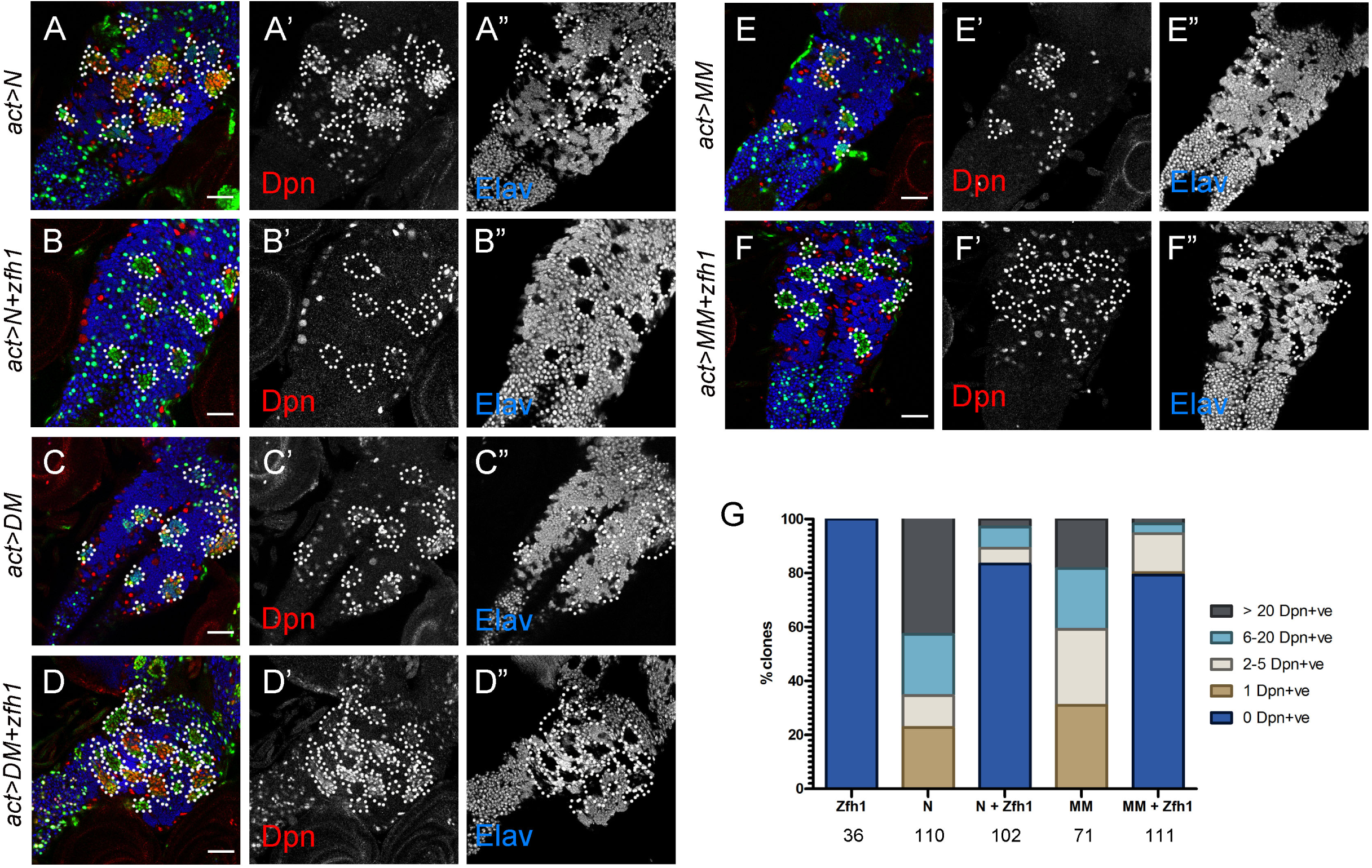
Effects of *zfh1* misexpression on *N, DM* and *MM* lineages. (A-E) FLPout (*act>STOP>Gal4*) clones (traced by dotted lines) expressing GFP (green) and the indicated transgenes. Dpn (red) marks NSCs and Elav (blue) marks neurons. (A’-E’) Dpn channel and (A”-E”) Elav channel in greyscale. All images are from VNCs. Scalebars 50μm. (G) 3d old (ACI) clones were scored for the number of NSCs per clone. The total number of Type I clones scored for each genotype is shown below the chart.

*gcm* overexpression had a markedly different effect in *N* vs *DM* tumour backgrounds (Figure 9, Figure 9 – supplement 1). In the N background it enhanced neuronal de-differentiation, with fully penetrant repression of Elav, but little or no repression of Pros (Figure 9 – supplement 1). Repo was dramatically induced and often co-expressed with Dpn (Figure 9). More specifically, N+gcm clones consisted mostly of two types of cells: (1) NSC-like Dpn-positive/ Repo-negative cells and (2) doubly Dpn & Repo-positive cells with lower Dpn levels. A third cell type, glia-like Dpn-negative/ Repo-positive cells, was also seen at lower numbers. In contrast, *gcm+Hes* co-expression did not induce Repo as strongly; only a few cells were Repo +ve in 10-20% of the clones, which is less than the frequency of Repo induction by misexpression of *gcm* alone. *DM/MM + gcm* also did not enhance Elav repression beyond that caused by *DM* or *MM* alone (Figure 9 – supplement 1B, compare with A, which is *N+gcm*). Instead, it reduced the NSC-like cells, as evidenced by a reduction in Ase- or Dpn-positive cells compared to *DM/MM* alone (Figure 9 – supplement 1D-H). In conclusion, whereas *gcm* synergizes with Notch to generate precursor-like cells, many of which express the glia marker Repo, this cannot be recapitulated by co-expressing *gcm* with Hes factors, suggesting a different mechanism of Notch-Gcm synergy that may not involve the usual Hes targets of Notch.

**Figure 9.**
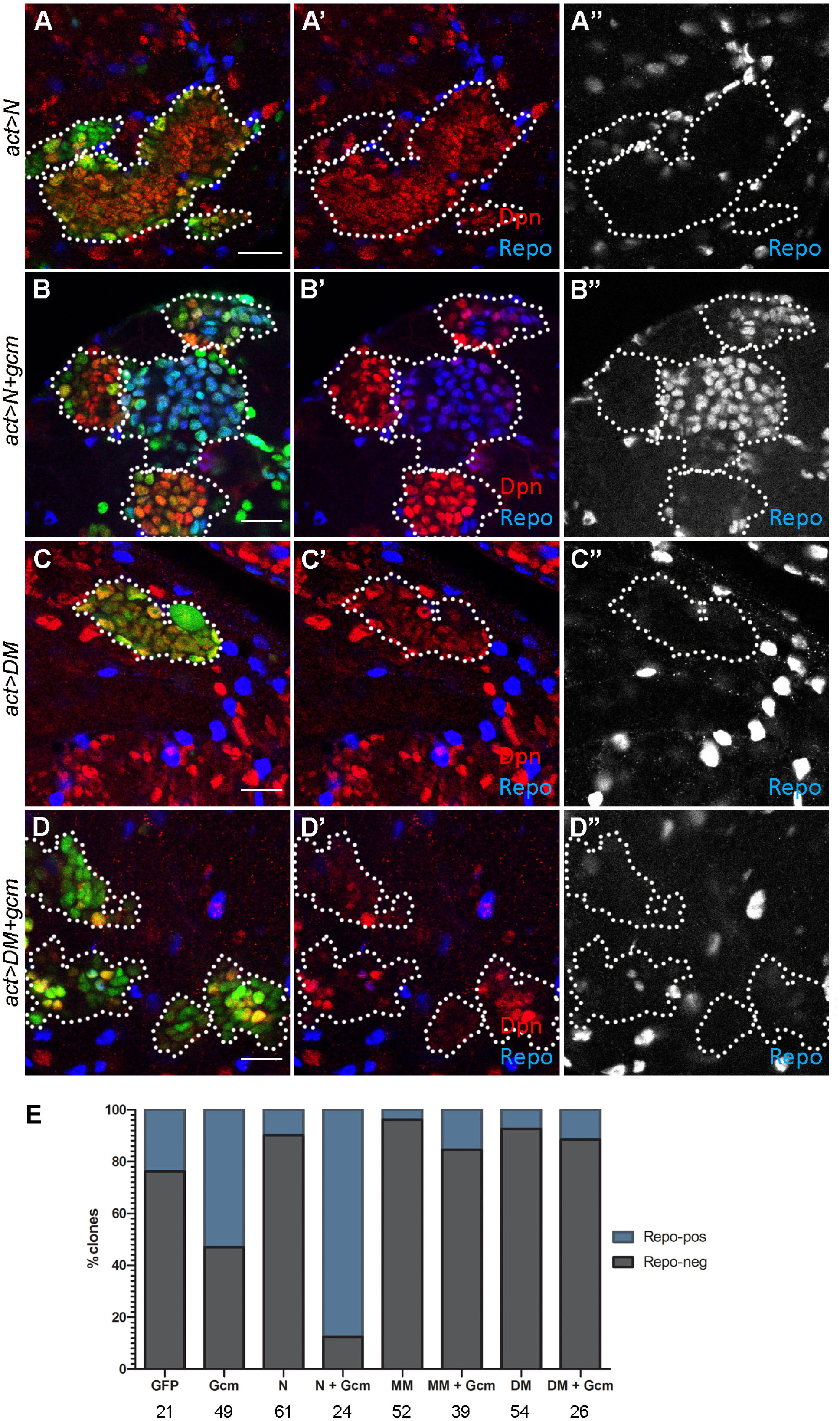
Effects of *gcm* misexpression on *N* and *DM* lineages. (A-D) FLPout (*act>STOP>Gal4*) clones (traced by dotted lines) expressing GFP (green) and the indicated transgenes. Dpn (red) marks NSCs and Repo (blue) marks glial cells. (A’-D’) shows an overlay of the Dpn and Repo channels; (A”-D”) shows the Repo channel only. Note the general scarcity of Repo positive cells within clones, with the exception of (B), where two clones contain large numbers of glia. Scalebars 20μm (E) The frequency of clones containing at least one Repo positive cell (blue) vs none (grey). The numbers below the chart indicate the absolute number of clones scored for each genotype. Only clones in central brain regions were scored.

To determine the malignancy potential of NSC tumours after coexpression of *zfh1* or *gcm* we transplanted brain lobes bearing *N* or *DM* clones into abdomens of adult females. Both *zfh1* and *gcm* significantly decreased the number of allografts generating malignancies; in one case, *DM+gcm*, no tumours could be propagated (Figure 10). In the other three cases, successful host colonization fell from 80-90% down to less than 50% and many of these tumours grew more slowly, taking 21-40 days to produce detectable GFP. Therefore, both *zfh1* and *gcm* have an anti-tumorigenic effect when co-expressed with *N* or *DM*. Upon histological examination of *N+zfh1* tumours, we observed a large number of Elav positive (neuron-like) cells, which were only sporadically seen in *N* tumours. Intriguingly when the malignant tumour from the original allograft (“T0”) was transplanted to a new host, the resulting tumour (T1) contained significantly fewer Elav positive cells and this was also true for T2, the next passage (Figure 10). Therefore, the *N+zfh1* CNS lineages have a greater tendency to differentiate after allografting, but they lose this ability upon serial passaging.

**Figure 10.**
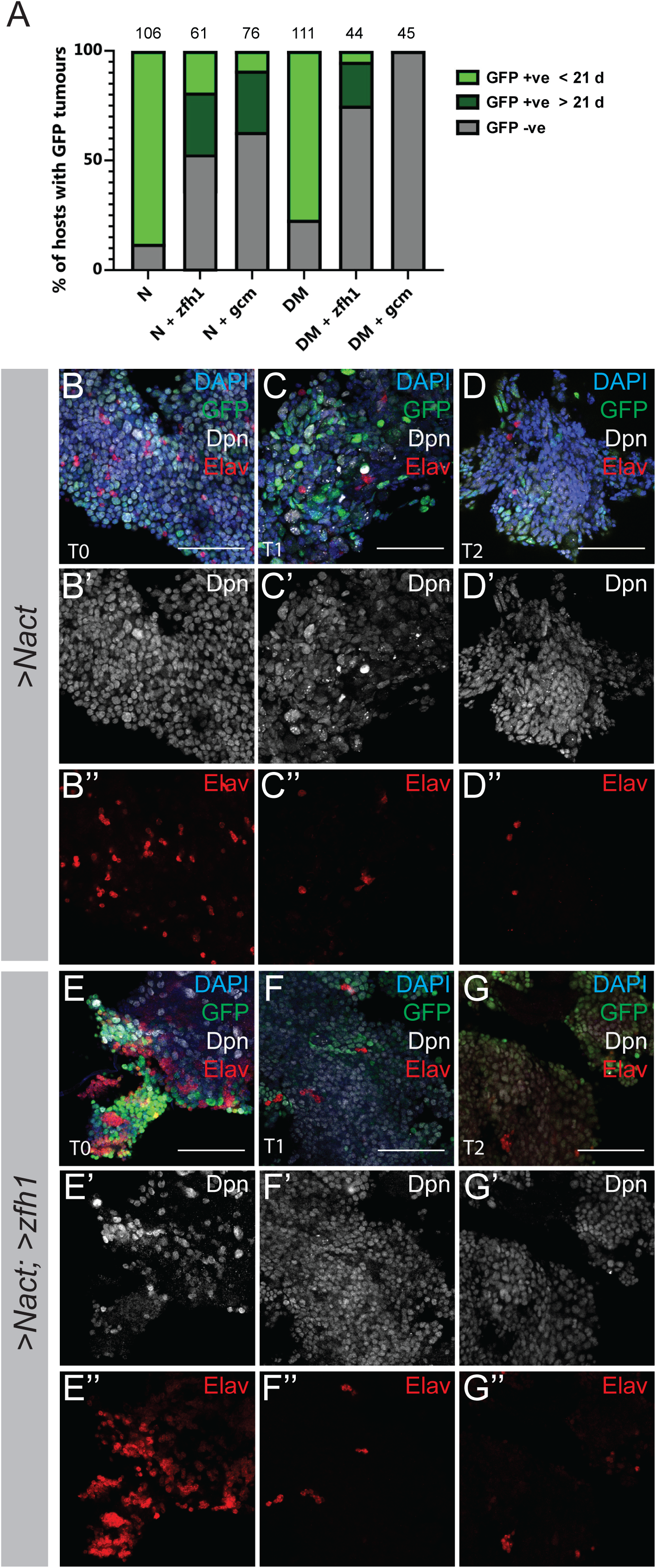
Effects of *gcm* and *zfh1* misexpression on *N* and *DM* tumorigenesis. A. Frequency of recovered tumours after transplantation of larval brain lobes carrying *act>STOP>Gal4* clones of the indicated genotypes. Tumours were assayed by detecting GFP in the transplanted host under the stereoscope. (B-D) Progression of *act>N* transplanted tumours. Shown are tumour fragments recovered from the host’s abdomen and stained for Dpn (white), GFP (green), Elav (red) and DAPI (blue). (B) T0 tumour recovered after transplanting larval brain lobe. (C) T1 tumour recovered after transplanting T0 tumour pieces. (D) T2 tumour recovered after transplanting T1 tumour pieces. (E-G) Progression of *act>N+zfh1* transplanted tumours stained as in B-D. Scalebars 25μm.

## DISCUSSION

During neurogenesis, NSCs divide asymmetrically and their more differentiated progeny sends a Dl-Notch signal back to the NSC. We have studied in detail the NSC hyperplasias caused by overactivation of Notch or overexpression of its *Hes* targets, *dpn, E(spl)mγ* and *E(spl)m8*, in the larval CNS of *Drosophila*. We have shown that such ectopic NSCs are highly prone to malignancy, as evidenced by their ability to be serially allografted and propagated in adult hosts. We have also performed a transcriptomic and chromatin occupancy analysis to identify target genes of Hes repressors that act as tumour suppressors in this context. The proliferating larval NSCs and their progeny were already known to be prone to tumorigenesis if their highly regulated mode of asymmetric cell divisions is perturbed. Shown initially for mutants in central factors that establish cytoplasmic and spindle asymmetry (Gateff 1978, Caussinus and Gonzalez 2005), tumorigenesis has been more recently documented for other insults, among others overexpression of the early temporal TF chinmo (Narbonne-Reveau et al. 2016) or loss of the differentiation TF nerfin1 (Froldi et al. 2015). We add the overactivity of the Notch-Hes axis to these initial insults that can lead NSC lineages to tumorigenesis.

How does the Notch-Hes axis contribute to NSC maintenance? Earlier we showed that Su(H), the TF that tethers cleaved (activated) Notch to its target enhancers, binds near all genes specifically expressed in the larval NSC, indicatively *klu, ase, wor, pnt, dm* and *mira*, besides the *Hes* genes in the *dpn* and *E(spl)* loci (Zacharioudaki et al. 2016). This is consistent with Notch signalling promoting NSC maintenance and with the fact that one of the steps taken by the differentiating NSC progeny is to downregulate the Notch signal by accumulating high levels of a Notch inhibitor, Numb. In fact the phenotype of our *act>>Gal4* (FLPout) clones overexpressing *NΔecd* or *2xHes* (Figure 1) is almost identical to the phenotype of *numb*^*-/-*^ clones, namely weak and variable hyperplasias in Type I lineages and extreme hyperplasias in Type II lineages (Lin et al. 2010).

From our earlier analysis, we knew that *Hes* genes are needed for NSC maintenance (Zacharioudaki et al. 2012). In the present work we have gained insight on how the Hes factors contribute to NSC maintenance. We have shown that Dpn, one of the Hes factors, binds and represses a number of transcription factor genes, among which are some that are thought to promote differentiation. A typical one is *gcm*, a well-known glia promoting TF, which was able to antagonize Notch/Hes-induced tumorigenesis (Figure 10) in addition to promoting gliogeneis (Figure 9). Another Hes-repressed direct target is *zfh1*. We showed that this TF represses *dpn* and *ase* (Figure 7). Previously, genetic interactions implicated Zfh1 in the action of the NURD chromatin remodelling complex, which was shown to shut off *E(spl)mγ* transcription in NSC progeny (Zacharioudaki et al. 2019). A plausible scenario is that Zfh1 acts with NURD to actively decommission many Notch-responsive target enhancers in NSC lineages, including *dpn* and *E(spl)*, the only Hes genes expressed in NSCs and needed for their self-renewal. Besides repressing *dpn* and *E(spl)* genes, Zfh1 must also have other NSC targets, since its forced expression ameliorates *DM* tumorigenesis, despite exogenous expression of *dpn* and *E(spl)mγ* (Figures 8 & 10). Likely candidates for other Zfh1 targets are the remaining NSC stemness TFs, like *ase, wor* and *klu*, although direct repression of other stem-cell functions, like the cell cycle, is also possible.

Other TFs reported to repress NSC markers are Pros (Vaessin et al. 1991, Choksi et al. 2006, Cabernard and Doe 2009, Southall and Brand 2009, Colonques et al. 2011, Liu et al. 2020), LolaN (Flybase: Lola-PP) (Southall et al. 2014) and Nerfin1 (Froldi et al. 2015, Vissers et al. 2018) in Type I lineages and Erm in Type II INPs (Weng et al. 2010, Janssens et al. 2014, Koe et al. 2014). All four, like Zfh1, show mutually repressive relationships with the Hes factors: they have strong Dpn binding in their vicinity (Figure 4) and most are downregulated in N/Hes tumours (Figure 3A, although *pros* downregulation was less than the 1.8x cutoff we imposed). Repression by Hes factors has been reported for *erm* (Janssens et al. 2017, Li et al. 2017) and *pros* (San Juan and Baonza 2011, Zhu et al. 2012) and we now show it for *zfh1* (Figures 7 & 8). Together Pros, Zfh1, LolaN and Nerfin1 may therefore constitute a group of TFs that switch off the expression of stemness markers, ensuring their specific accumulation only in the NSCs.

Interestingly, these anti-stemness TF genes (*zfh1, erm, nerfin1, lola* and *pros*) are also in the vicinity of experimentally detected Su(H) binding sites, even though Notch activity represses them (Zacharioudaki et al. 2016) and this study). Perhaps high Notch signalling in the NSC primes these chromatin regions for activity, but transcription is prevented by the binding of Hes repressors that accumulate to high levels in the NSC. As Notch signalling diminishes in the NSC progeny, Hes proteins disappear and these genes are derepressed. They in turn would serve to repress residual Hes transcription (and that of other stemness genes) in the early-born neurons, where lower levels of Notch activity are still present, as evidenced by the need for Notch in neuron fate decisions (Truman et al. 2010) and the neuronal expression of *Hey*, another Notch target (Monastirioti et al. 2010). This mutual repression between stemness and anti-stemness factors may be a way to ensure the gradual and orderly differentiation of neurons during the 6 days of post-embryonic neurogenesis, so that they have time to generate the complex neuronal circuits of the adult animal.

Do these anti-stemness factors directly promote differentiation, like Gcm? We have evidence that Zfh1 does the contrary, namely antagonizes differentiation. We see this by the loss of early neuronal markers like Elav upon Zfh1 overexpression, especially upon its co-overexpression with NΔEcd or Hes factors (Figure 8). A plausible scenario is that after GMC divisions, newly-born neurons remain in a developmentally plastic state by virtue of expressing *zfh1* (and probably also *pros, lolaN* and *nerfin1*) and they only differentiate at a later time, perhaps in response to developmental cues arising during animal maturation. When in this plastic state, cells can be reprogrammed by increased Notch/Hes activity and may even re-enter the cell cycle, leading to ectopic NSCs. Note that in our experiments mature neurons and glia never de-differentiated to NSCs upon the same Notch/Hes stimulation. A similar role in delaying differentiation has been described for Zfh1 and its mammalian homologue ZEB1 during myogenesis (Siles et al. 2013, Boukhatmi and Bray 2018). We also noted a difference between the activity of Zfh1 and mammalian ZEB1: Whereas ZEB1 is well-known for promoting epithelial-to-mesenchymal transition and enabling tumour progression and metastasis (Brabletz and Brabletz 2010, Rosmaninho et al. 2018), in our system, Zfh1 suppressed NSC lineage tumorigenesis.

In summary, Notch activity favours NSC fate via setting up a network of TFs that mutually cross-regulate and allow a gradual transition to differentiation as Notch activity gets turned down. Why does persistent Notch activity lead to malignant tumours? Our data show that tumorigenesis is not a simple consequence of inability to differentiate. *N/zfh* and *DM/zfh* CNSs contain more undifferentiated cells than *N* or *DM*, but they are also less tumorigenic. Also, normal NSCs, despite being undifferentiated, do not produce tumours when FACS-sorted and allografted (Landskron et al. 2018). Further studies will be needed to characterize the transition of NSC lineages to malignancy.

## MATERIALS AND METHODS

### DNA constructs

UAS constructs of *dpn, E(spl)mγ* and *E(spl)m8* were made by PCR amplifying the coding sequences of these genes (primer sequences are listed in supplementary materials and methods) from a cDNA template. The cDNA was synthesized from RNA extracted from 30 adult *yw* flies. PCR fragments were subsequently cloned into the pENTR3C entry vector (http://www.ciwemb.edu/labs/murphy/Gateway%20vectors.html) and subsequently N-terminally tagged with 3xHA epitope using the UASt destination vector pTHW by Gateway cloning technology.

These plasmids were injected along with helper plasmid (Δ2–3 –transposase) (Karess and Rubin 1984) into embryos of *yw* to generate transgenic *UAS-HA-dpn, UAS-HA-E(spl)mγ* and *UAS-HA-E(spl)m8* flies.

### Fly stocks and genetics

*Drosophila* stocks are described in Flybase and were obtained from the Bloomington Stock Center unless otherwise indicated. *zfh1-lacZ* is BDSC stock 11515 that carries the *P {PZ}zfh1*^*00865*^ enhancer trap. *gcm-lacZ* is BDSC stock 5445, that carries the *P{PZ}gcm*^*rA87*^ enhancer trap.

Overproliferating third instar larval CNSs were obtained by crossing *UAS-HA-mγ* and/or *UAS-HA-dpn*, flies with *tubP-Gal80ts; UAS-CD8GFP; grhNB-Gal4* (Zacharioudaki et al. 2016). Crosses were maintained at 18°C for 8 days, then transferred to 30°C for 24hrs prior to dissection.

#### Flip-out clones

Stocks for act-flip-out clones were generated as following: Initially the act-FRT>STOP>FRT-Gal4 (BDSC stock 4780) was recombined with UAS-nlsGFP and subsequently combined with hs-FLP. Flies from the hs-FLP; act-FRT>STOP>FRT-Gal4, UAS-GFP stock were crossed with the appropriate *UAS* combinations for generating clones. *UAS-zfh1* is BDSC stock 6879 and expresses the long isoform of *zfh1* (RB). *UAS-gcm* was obtained from Fly ORF (Bischof et al. 2013). Control clones were obtained by replacing UAS-lines with *yw*. Progeny underwent heat shock for 45min-1h at 37°C at 72h after egg lay (AEL). Phenotypes were analysed 3 days later, unless otherwise indicated.

#### For mosaic analysis (MARCM)

*FRT82 zfh1*^*75*.*26*^ (Leatherman and Dinardo 2008), *FRTG13 gcm*^*N7-4*^ (Bernardoni et al. 1997) were crossed to appropriate *FRT tubP-Gal80* counter-chromosomes (Lee and Luo 1999) combined with *hs-FLP, tubP-Gal4, UAS-GFP* for generating clones. Progeny underwent heat shock for 1h at 37°C at 72h AEL and CNSs were dissected out from wandering 3^rd^ instar larvae 3 days post clone induction.

### Expression arrays

For each expression array, RNA was isolated from 300 larval CNSs dissected in cold PBS using the RNeasy Plus Mini Kit (Qiagen). Synthesis of double-stranded cDNA and biotin-labelled cRNA was performed according to manufacturer’s instructions (Affymetrix). Fragmented cRNA preparations were hybridized to *Drosophila* genome oligonucleotide arrays (GeneChip Drosophila Genome 2.0 Array (3’ IVT Expression Affymetrix)] using an Affymetrix hybridization Oven 640, washed, and then scanned on a GeneChip Scanner 3000.Three replicate arrays were analyzed for each genotype. Data from expression arrays have been deposited to Gene Expression Omnibus with accession number GSE141794.

Gene expression values were obtained using Affymetrix Transcription Analysis Console from CEL intensity files using MAS5 without normalization and further quantile-normalized and then filtered for low expression. Differential gene expression analysis was performed on a set of total 8479 probesets with detectable expression. For each of the four mutant conditions log_2_-fold-change values were calculated against wild-type samples, coupled with a Benjamini-Hochberg adjusted p-value of a two-tailed t-test.

Differentially expressed genes were defined in the reduced set, based on moderately conservative criteria of absolute log_2_(fold-change) greater or equal to 0.85 (corresponding to fold-change values of >1.8 or <0.55) and corrected p-values of less or equal to 0.05 (FDR 5%). The union of all genes differentially expressed in at least one of the studied conditions comprised 1410 genes. These were further clustered in 6 groups depending on their differential expression across conditions: Over-expressed in Notch (114 genes), Hes (417) or both Notch and Hes (42), under-expressed in Notch (242), Hes (484), or both Notch and Hes (96). Further comparison of transcriptomes was performed through Pearson correlation calculations of differential expression values for the set of the 1410 inclusive differentially expressed genes.

Comparison of differential expression with public datasets was performed through the application of simple thresholds corresponding to the top/bottom 10% of genes being differentially expressed in FACS-sorted neural stem cells vs differentiated neurons (Berger et al. 2012). Differential expression values in the four studied conditions were then analyzed for sets of ∼1600 genes corresponding to neuron and neuroblast-specific expression respectively (Figure 3 – supplement 2). To compare our CNS transcriptomes with those of other CNS tumours, we obtained data for *l(3)mbt* (Janic et al. 2010), *l(2)gl, scrib* (Richardson lab, GEO:<u>GSE48852</u>) and two *brat* RNAi genotypes (Neumuller et al. 2011) and cross-analysed gene expression levels for both our restricted 1410 differentially expressed genes (Figure 3 – supplement 3), as well as against the extended set of >8000 measured genes (Figure 3D). Clustering of gene expression profiles was performed on the basis of Pearson Correlation Coefficient similarity for all 8479 genes.

We used PANTHER (http://www.pantherdb.org/) (Mi et al. 2019) to assess statistical overrepresentation. Genes were provided as lists of Flybase FBgn IDs (Thurmond et al. 2019) to PANTHER and analyzed at the levels of GO (Biological Process and Molecular Function). An adjusted p-value of ≤ 0.05 was set as threshold for significance.

### Quantitative PCR (qPCR)

Larval CNSs of different genotypes were dissected and kept in Trizol where they were homogenized with a plastic pestle and incubated for 10 min on ice. RNA was extracted using chloroform and precipitated in isopropanol overnight at -20°C. RNA was resuspended in RNase free water. 1μg of RNA was reverse transcribed with oligo(dT)_15_ Primers (Promega C1101)] using SuperScript III Reverse Transcriptase (Invitrogen, now Thermo Fisher Scientific 18080044) Treatment with RNAseH removed RNA. The cDNA products were subsequently diluted 1:5 and 2μl were used as a template in each quantitative PCR reaction. Quantitative PCR was performed using KAPA SYBR Green fast Master PCR Kit (Roche SFUKB-KK (04707516001). Generation of specific PCR products was confirmed by melting-curve analysis. Ct values for all genes were normalized to the levels of *RpL32*. For data analysis, the delta-delta Ct values was applied. Primer sequences are listed in the supplementary Materials and Methods.

### Chromatin Immunoprecipitation

For each ChIP, chromatin was prepared from 500 third instar larval CNSs dissected from cross grhNB-Gal4> UAS-mγ; UAS-dpn. Tissues were cross-linked with 1%FA, quenched with 200mM Glycine. After homogenization in Nuclear Lysis Buffer (50 mM Tris-HCl pH 8, 10 mM EDTA, 1% SDS and Protease inhibitor), chromatin was fragmented using Diagenode Bioruptor for 7 cycles with 30sec OFF/30sec ON to obtain 200-500bp length. Fragmented chromatin was immunoprecipitated with anti-Dpn antibody (see Supplementary Materials & Methods) overnight at 4°C. Complexes were incubated with Protein A/G agarose beads (SC-2003) and precipitated chromatin was eluted using IP Elution Buffer (IPEB) (100 mM NaHC03, 1% SDS) and purified using Qiagen PCR purification kit.

Validation of enrichment was performed over specific chromatin regions (Primer sequences in supplementary material and methods).

### Ion Torrent library sequencing and data processing

Two biological replicates of anti-Dpn ChIP and one input were sequenced on Ion Torrent Proton platform with Ion 540 ChIP kits. Libraries were prepared according to manufacturer’s protocol using the Ion Plus Library Kit (# 4471252).

Fastq reads were mapped to UCSC/dm3 genome (archive-2015-07-17-14-30-40) downloaded from iGenomes using bowtie2 (version 2.2.8, --very-sensitive) (Langmead and Salzberg 2012). Bedgraphs of mapped reads were generated by bedtools genomecov (v2.25.0) and uploaded on UCSC/BDGP R5/dm3) genome browser for visualization. Reads that overlapped with dm3 repeat elements (downloaded from UCSC,) were removed from bam files prior to peak calling using bamtools intersect (default). Peak calling was performed with macs2 over input (-p 0.05, version 2.1.0.20140616) (Zhang et al. 2008). A Dpn binding consensus list of 229 genomic regions was generated using peaks with FC>2 over input in both biological replicates. Motif enrichment analysis was performed in cistrome (Liu et al. 2011) using the SeqPos motif tool, searching against known *Drosophila melanogaster* transcription factor binding motifs from Jaspar. Genome wide peak distribution was performed with the pavis tool (Huang et al. 2013) using the flybase R5.57/dm3 genome assembly and a 5kb upstream and 5kb downstream length window. Peak to gene annotation was performed by bedtools intersect using the Dpn binding consensus and the RefseqFlat.txt with 5kb upstream and 5kb downstream window extensions for each Refseq transcript. Gene ontology analysis of the genes with peaks in their proximity was performed using flymine v47.1 (Lyne et al. 2007). ChIP data have been deposited in Gene Expression Omnibus (GSE141794).

### Immunohistochemistry

Fixation and immunohistochemistry of larval tissues were performed according to standard protocols. Details of primary antibodies are provided in the supplementary Materials and Methods. Mouse, rabbit, guinea pig or rat secondary antibodies were conjugated to Alexa 488, 555, 568, 633 or 647 (Molecular Probes, now Thermo Fisher) or to FITC, Cy3 or Cy5 (Jackson ImmunoResearch). Samples were imaged on a Leica SP8 confocal microscope (Confocal Facility, IMBB, FORTH).

### Transplantation assay

Transplantation assays were performed as previously described by Rossi and Gonzalez (Rossi and Gonzalez 2015). Donor larval brains bearing FLPout clones overexpressing different transgene combinations were dissected, sliced, loaded into a fine glass needle and implanted into the abdomen of female host w1118 flies using a nanoinjector (Nanoject II Auto-Nanoliter Injector, Drummond Scientific Company, 3-000-205A). Host flies carrying allografts were kept at 25°C and examined daily for the presence of GFP in their abdomen and other tissues. Malignant GFP positive tumor pieces (T0) were dissected out of the abdomen of host flies and either re-transplanted into new host flies (T1) or fixed with 4% formaldehyde (for 25 min at RT) and used for immunohistochemistry experiments according to standard protocols.

## Supporting information

Magadi et al Supplement

## FIGURE LEGENDS

**Figure 1-Suppl1 *E(spl)* gene expression in the brain lobe**. Each of the seven bHLH E(spl) proteins (m8, m7, m5, m3, mβ, mγ, mδ) was visualized as a GFP fusion from a transgenic genomic BAC transgene (Kudron et al. 2018). E(spl)-GFP is green, Dpn (NSCs) blue and Elav (neurons) red in all panels. In the central brain, white arrowheads mark NSCs; yellow arrowheads mark GMCs/early neurons; magenta arrowheads mark glia. In the optic lobe, white arrows mark medulla NSCs, yellow arrows mark GMCs/early neurons and magenta arrows mark other structures, like the very prominent neuroepithelium. All panels show ventral superficial sections, except E, which shows a deep cortical section to highlight neuropil glia expression of E(spl)m7. Scalebars 50μm.

**Figure 1-Suppl2 Notch and Hes induced hyperplasia in Type II lineages**. Panels A-D are stained to reveal NSCs (Dpn, red) and GMCs/neurons (Pros, blue). FLP-out clones are marked with GFP (green) and overexpress (A) GFP alone (control); (B) UAS-m8, UAS-mγ (MM); (C) UAS-dpn, UAS-mγ (*DM*); (D) UAS-NΔecd (N). Clones are outlined. In panels A and B adjacent smaller Type I clones are also outlined and marked with an arrow. (E) Cartoons depict the cellular composition of wt vs *N* overexpressing TypeII lineages. *MM* and *DM* lineages would be the same, only they should additionally have a few blue cells (neurons). Scalebars 20μm.

**Figure 1-Suppl3 Further examples of neural hyperplasia caused by N/Hes overactivity**. All panels show *act>STOP>Gal4* clones expressing GFP and transgenes as marked. A-D: NSCs/GMCs are detected by Ase (red); GMCs/young neurons are detected by Pros (blue) – individual channels are shown in greyscale. A is essentially wt, as *dpn* expression alone rarely causes weak defects in Type I lineages. White arrows mark examples of Ase positive/ Pros negative (NSC-like) cells. Yellow arrows mark double-positive (GMC-like) cells. E: Dpn (red) marks NSC-like cells (examples shown by white arrows) and Pros (blue) marks GMC/neuron-like cells. Examples of doubly Dpn/Pros positive cells are shown by magenta arrows. Scalebars 20μm.

**Figure 1-suppl4 N/Hes NSC hyperplasias persist after pupariation**. A,B: brain lobes dissected at 24h APF; C,D: brain lobes dissected from freshly eclosed adult escapers. All animals expressed GFP together with the indicated transgenes from an act>STOP>Gal4 driver after hsFLP induction at early larval stages. Dpn is stained red and Pros blue. Individual channels shown in greyscale. The two clones that retain a Dpn-positive NSC in the wt (A) are mushroom body lineages that continue proliferating into the early pupal stages. Scalebars 50μm.

**Figure 3-Suppl1 qPCR validation of select mRNAs**. Expression levels were calculated by q-RT-PCR from *DM* or *MM* overexpressing CNSs (using *grhNB-Gal4*) or wt controls. Expression levels are shown relative to *RpL32* RNA. Error bars show the standard error of the mean from triplicate measurements. Asterisks indicate samples that are significantly different from control by Student’s t-test (* P<0.05, **P<0.01).

**Figure 3-Suppl2 Comparison of N/Hes differentially regulated genes with neuron, GMC and NSC-enriched gene-sets. (A**,**B)** Comparison with the neuron vs NSC enriched genes defined by Berger et al (2012): they show differential enrichment in FACS-sorted neuron or NSC populations from wt CNSs. (A) Heatmaps showing the log_2_(fold-change) of neuron-enriched genes (left) and NSC-enriched genes (right) in the N and Hes overexpression backgrounds. (B) Venn diagram showing overlap of high confidence (FDR≤ 0.05) UP or DOWN-regulated genes in the N/Hes conditions (union of *N* with *DM* and *MM*, as shown in Figure 3B,C) with the neuron vs NSC enriched gene-sets. (C, D) Comparison with the NSC vs GMC enriched genes defined by Wissel et al (2018): dissociated NSCs were cultured in vitro and their smaller GMC progeny FACS-sorted from the larger NSCs. 3 and 5 refer to the hours of culturing. (C) Heatmaps showing the log_2_(fold-change) of GMC-enriched genes and NSC-enriched genes from each culture timepoint in the N and Hes overexpression backgrounds. (D) Venn diagram showing overlap of high confidence (FDR≤ 0.05) UP or DOWN-regulated genes in the N/Hes conditions (as in B) with the GMC vs NSC enriched gene-sets from the two culture timepoints.

**Figure 3-Suppl3 Comparison of N and Hes CNS tumours with other CNS tumours. A**. Heat map of the fold-change of the 1410 genes selected by differential expression in the N or Hes tumours (Figure 3A) in five additional tumour transcriptomes. KK and GD are two *brat* RNAi genotypes (Neumueller et al Cell Stem Cell 2011), whereas scrib, lgl and mbt refer to homozygous mutant larval CNSs for the genes *scrib, l(2)gl* and *l(3)mbt*, respectively, all of which show CNS hyperplasia.

**Figure 5-Suppl1 *zfh1-lacZ* expression in the wt**. β-galactosidase is green in all panels. (A) Brain lobes counterstained with glial marker Repo (blue) and NSC marker Dpn (red). (B-D) Counterstaining for neuronal marker Elav (blue) and GMC/young neuron marker Pros (red) – note that Pros is also expressed in some glial cells. (B) Brain lobes. (C-D) VNC, ventral side (C) and dorsal side (D). Examples of GMCs/early neurons (Pros and Elav positive) are shown with yellow arrowheads, mature neurons (Elav positive/ Pros negative) by white arrowheads and glia (Repo positive) by magenta arrowheads. Scalebars 50μm

**Figure 6-Suppl1 Loss-of-function analysis of *zfh1* and *gcm***. MARCM clones are marked with GFP (green) and traced by dashed outlines. (a) WT, (B) clones are homozygous for *gcm*^*N7-4*^ (C) clones are homozygous for *zfh1*^*75*.*26*^. NSCs are imaged by Dpn (red) and GMCs/young neurons by nuclear Pros (blue). Individual channels shown in greyscale. Scalebars 20μm.

**Figure 7-Suppl1 Effect of *zfh1* misexpression on CNS lineages**. *act>STOP>Gal4* FLPout clones are marked by GFP (green) and traced with dotted outlines. (A) wt clones 3d ACI stained for Pros (blue, shown separately in A’). (B) *UAS-zfh1* clones 2d ACI stained for Pros (blue, shown separately in B’). (C,D) *UAS-zfh1* clones 1d (C) or 2d (D) ACI stained for Ase (red) shown separately in C’, D’. (E) wt clones 1d ACI stained for Dpn (red) and the mitotic marker phosphor-H3 (blue). (F) *UAS-zfh1* clones 1d ACI stained as in E. (G,H) Examples of brain lobes carrying UAS-zfh1 clones 3d ACI and stained for the glia marker Repo (blue) and Dpn (red) in G or Ase (red) in H. G is a more superficial section than H. A-D scalebars 10μm, E scalebar 15μm, F scalebar 10μm, G-H scalebars 15μm.

**Figure 7-suppl2 Effect of *gcm* misexpression on CNS lineages**. *act>STOP>Gal4* FLPout clones are marked by GFP (green) and traced with dotted outlines. (A,C,E) Control clones (GFP only); (B,D,F) UAS-gcm clones. (A,B) Sample stained for Dpn (NSCs, red) and Elav (neurons, blue). (C,D) Sample stained for Dpn (NSCs, red) and Pros (GMCs/ young neurons, blue); note two Type II clones in C (arrows). (E,F) Sample stained for Dpn (NSCs, red) and Repo (glia, blue). In E and F deep sections are shown where clone cells may have migrated away from the superficial stem cell (superficial lineages marked by arrows in E; note that the upper one derives from the OL); nevertheless in E there is hardly any GFP marking of glia, whereas in F most clonal cells have adopted a glial fate. All scalebars 50μm.

**Figure 8-suppl1 Effect of *zfh1* misexpression on *N, DM* and *MM* lineages**. *act>STOP>Gal4* FLPout clones are marked by GFP (green) and traced with dotted outlines. (A-D) Effects on Ase (blue) and Pros (red). Most Type I N/Hes clones lose Ase upon zfh1 expression, with some exceptions (arrow in D). (E-H) Effects on Repo. Even though there are several Repo positive cells adjacent to clones (red arrows), there are very few within clones (green arrow in H’). In E-H blue is Dpn. Scalebars 25μm.

**Figure 9-suppl1 Effect of *gcm* misexpression on *N, DM* and *MM* lineages**. (A-E) *act>STOP>Gal4* FLPout clones are marked by GFP (green) and traced with dotted outlines. (A,C) misexpression of *NΔecd+gcm*. (B,E) misexpression of *DM+gcm*, (D) misexpression of *DM* (control for E). (F-H) Effect of *gcm* on the induction of Dpn (blue) by MM. F has no lineage clones (so the NSC number is not affected), whereas G and H have a large number of clones (see GFP-only channel in F”-H”). (I) Clones of the indicated genotypes were scored for the number of NSCs per clone. The total number of Type I clones scored for each genotype is shown below the chart. Scalebars: A-E 25μm, F-H 50μm.

### ACKNOWLEDGEMENTS

Funding for this work was provided by the EU-Marie Curie Sklodowska action HEALING (ITN consortium), a Worldwide Cancer Research grant to CD, a Fondation Sante grant to CD, a Hellenic Foundation for Research & Innovation grant to EZ and two short term grants, EMBO and Company of Biologists, to SSM. We are grateful to Sarah J. Bray and Cayetano Gonzalez for hosting SSM and training him in chromatin immunoprecipitation and tumour allografting. We thank Ioannis Livadaras and Alexandros Babaratsas for fly injections and maintenance. Microarray hybridizations and high throughput sequencing were performed at the IMBB genomics facility. Many thanks to Erika Bach, Angela Giangrande, Yuh Nung Jan, James Skeath, Ruth Lehmann, Stephen DiNardo and Isabel Guerrero for providing fly stocks and antibodies.

## Supplementary Materials and Methods

**Primers list: cloning**

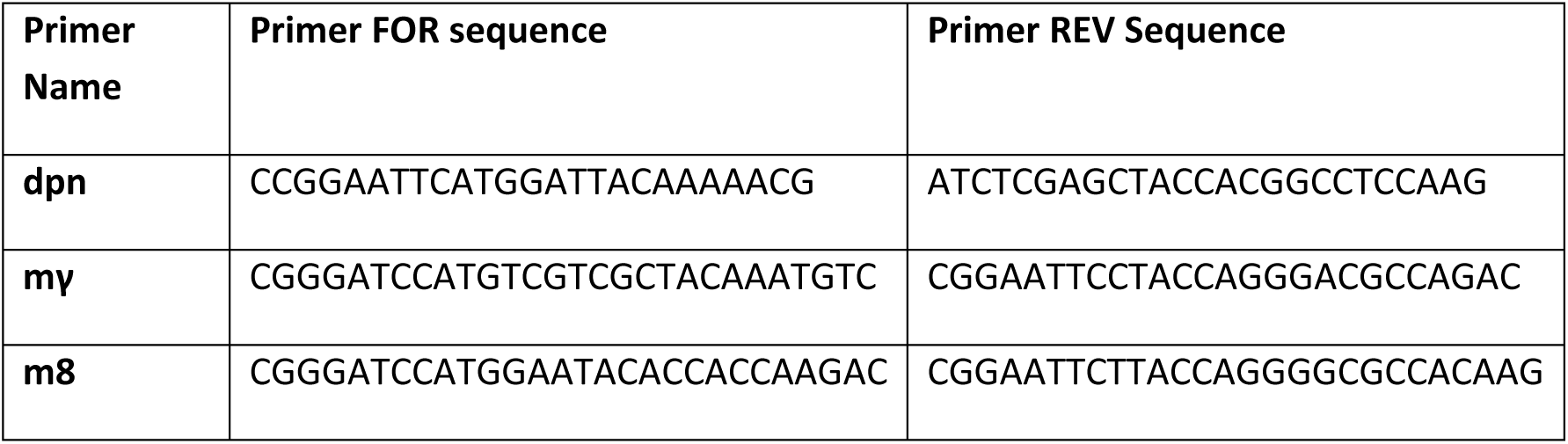

**Primers list: qRT-PCR**

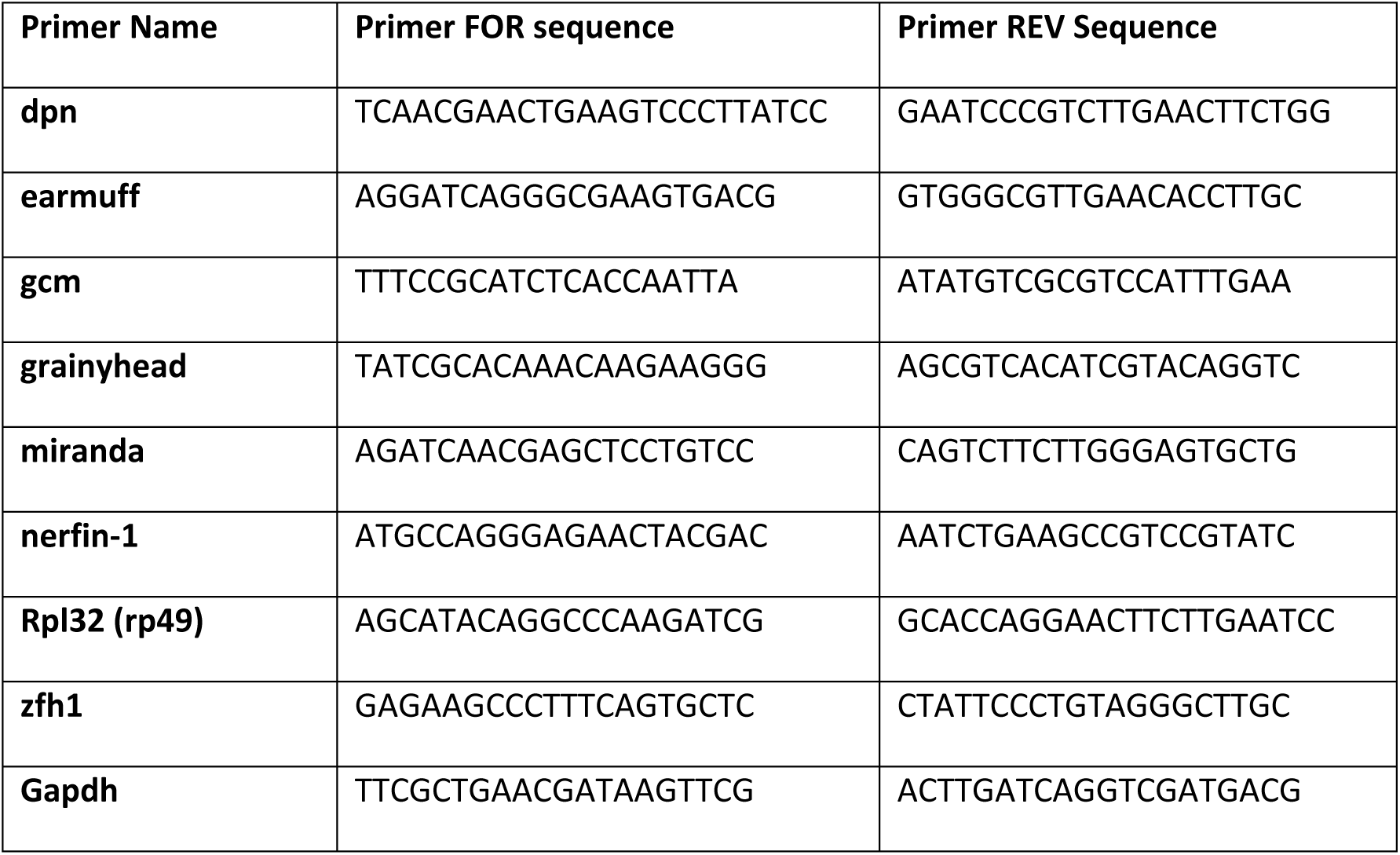

**Primers list: ChIP**

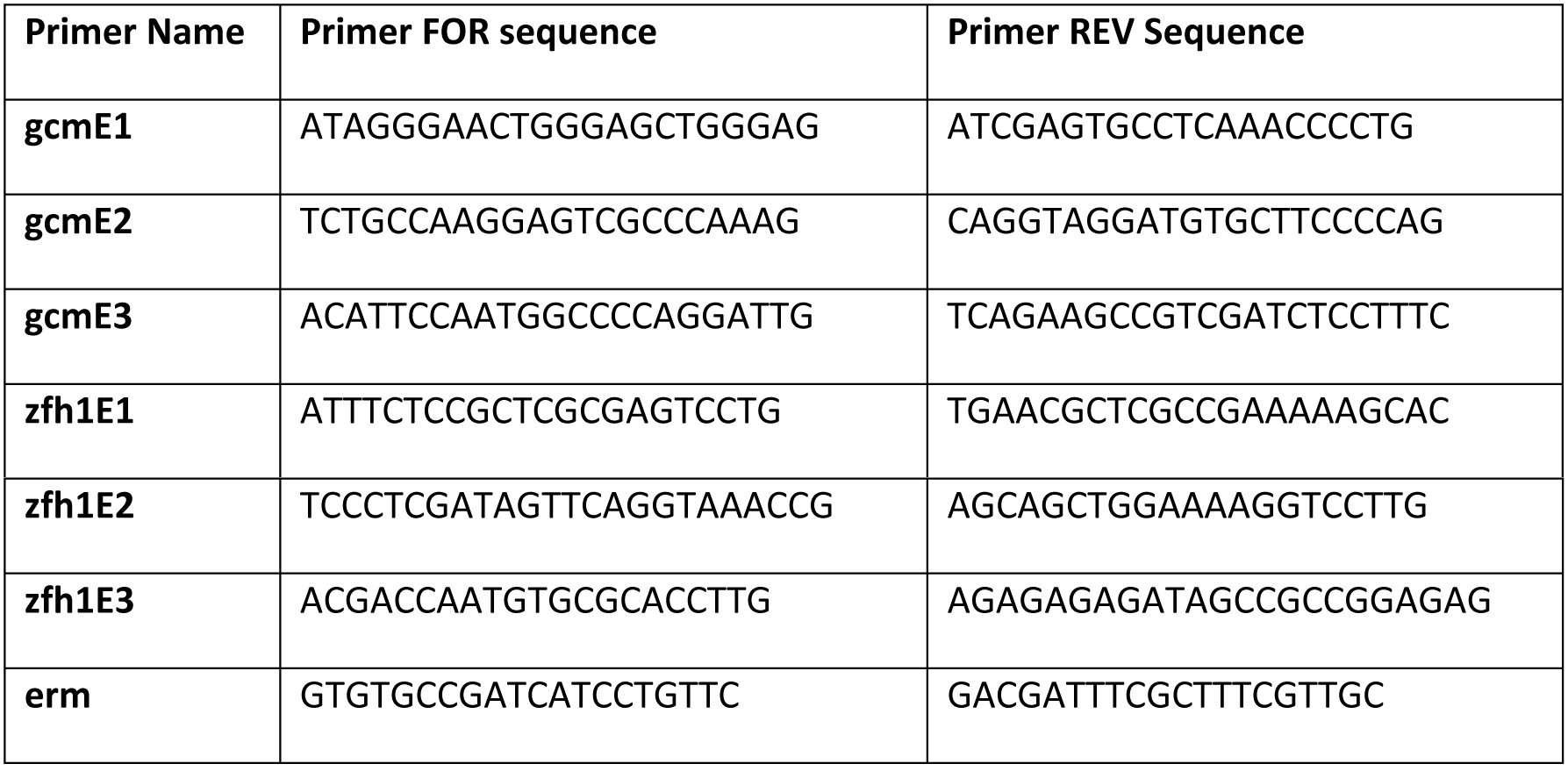

### Deadpan antibody production

Dpn ORF cloned in pET29a vector was a gift from James Skeath (Skeath et al. 2017). Protein production was performed according to Novagen manual for protein expression. Ni-NTA beads (Qiagen cat# 30210) purified protein was run on SDS-PAGE, the protein bands were excised and sent to Davids biotechnologie for guinea-pig immunization. The serum obtained caprylic acid treated, dialysed against 1x PBS and aliquots were stored at -80°C.

### List of 1° antibodies

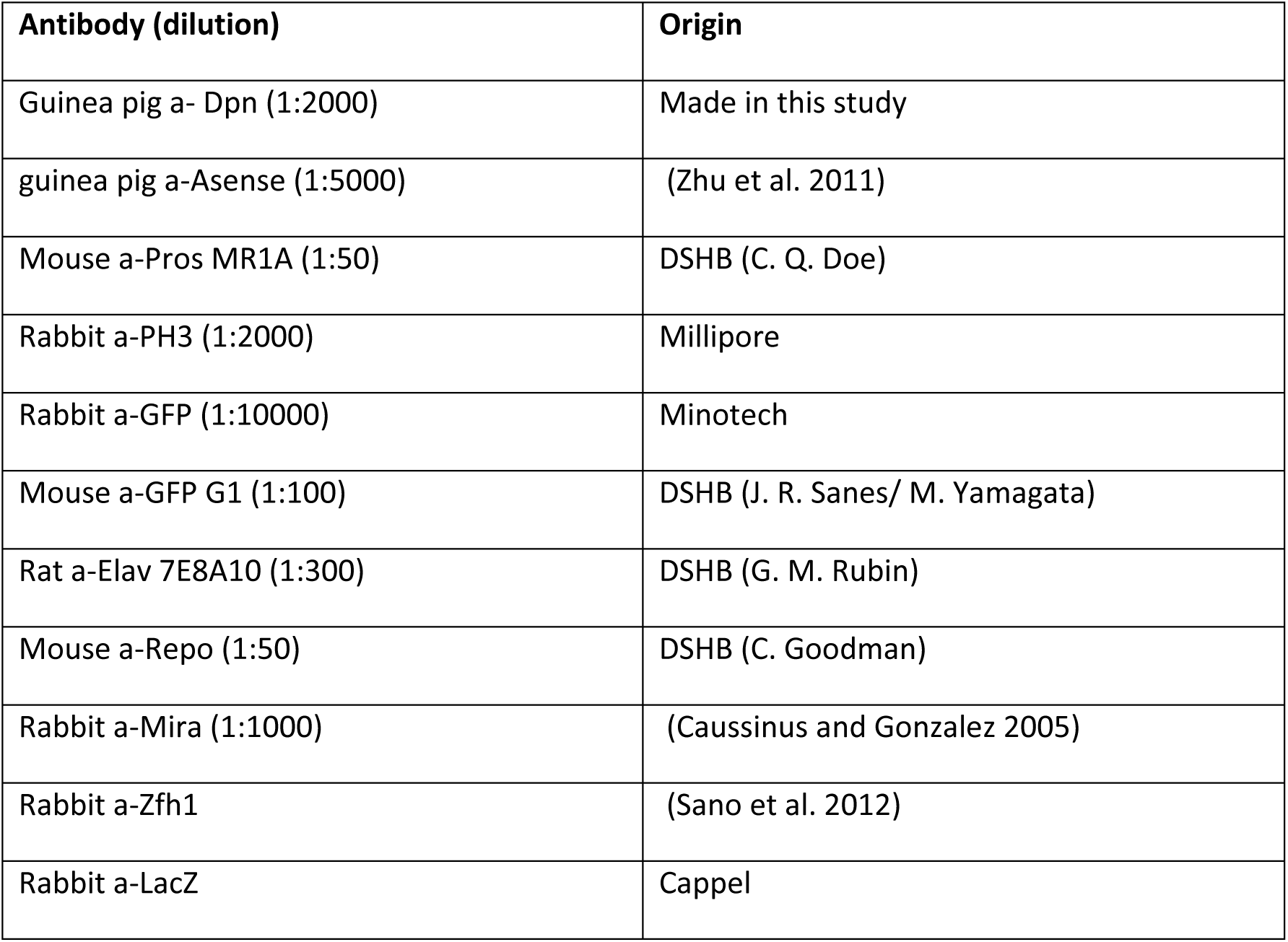

The monoclonal antibodies marked “DSHB” were obtained from the Developmental Studies Hybridoma Bank, created by the NICHD of the NIH and maintained at The University of Iowa, Department of Biology, Iowa City, IA 52242; in parenthesis we list the investigator who developed each mAb.

## Notes

https://www.ncbi.nlm.nih.gov/geo/query/acc.cgi?acc=GSE141794

