## Supplementary material for "Dissecting Hes-centered transcriptional networks in neural stem cell maintenance and tumorigenesis in *Drosophila*": Magadi et al Supplement

### Figure 1 - Supplement 1

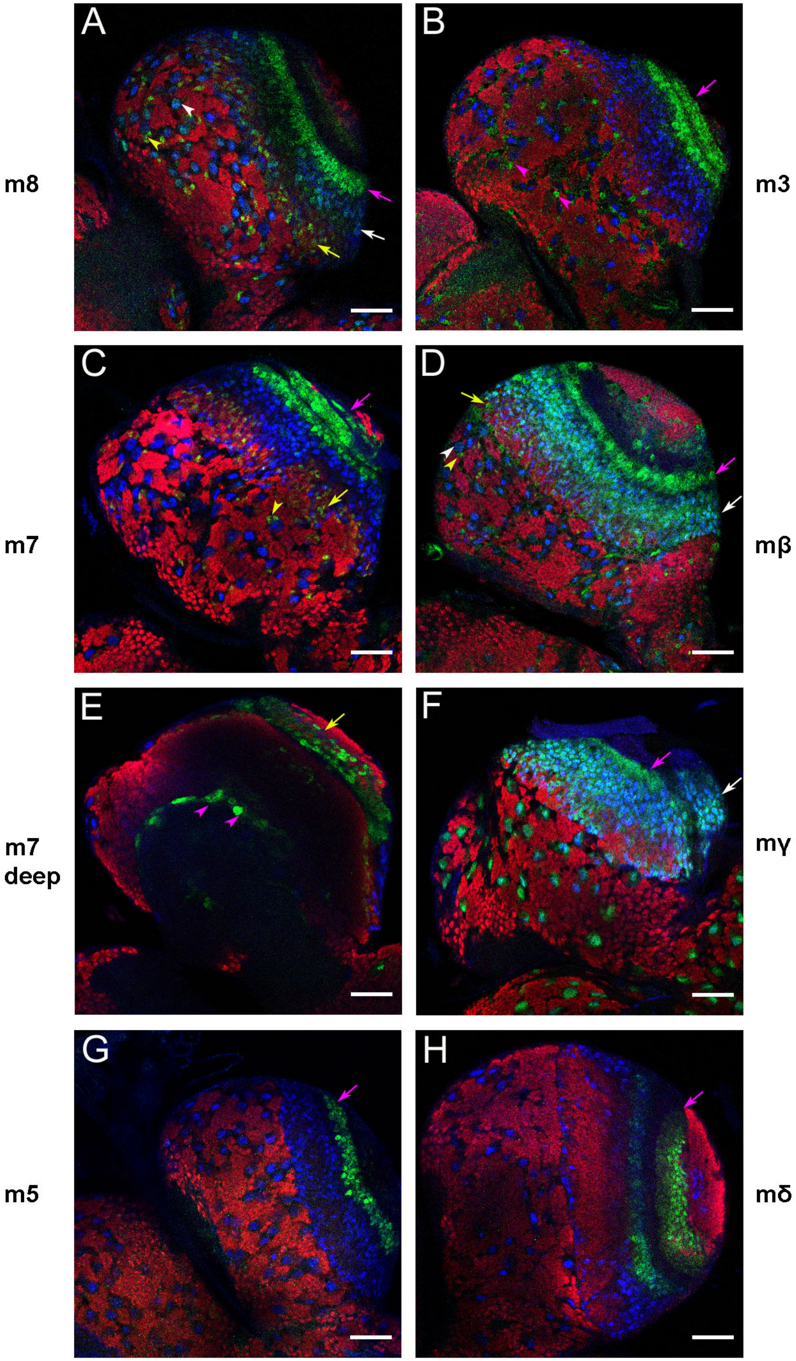

**Figure 1-Suppl1 *E(spl)* gene expression in the brain lobe.** Each of the seven bHLH E(spl) proteins (m8, m7, m5, m3, mβ, mγ, mδ) was visualized as a GFP fusion from a transgenic genomic BAC transgene{Kudron, 2018 #2231}. E(spl)-GFP is green, Dpn (NSCs) blue and Elav (neurons) red in all panels. In the central brain, white arrowheads mark NSCs; yellow arrowheads mark GMCs/early neurons; magenta arrowheads mark glia. In the optic lobe, white arrows mark medulla NSCs, yellow arrows mark GMCs/early neurons and magenta arrows mark other structures, like the very prominent neuroepithelium. All panels show ventral superficial sections, except E, which shows a deep cortical section to highlight neuropil glia expression of E(spl)m7. Scalebars 50µm.

### Figure 1 - Supplement 2

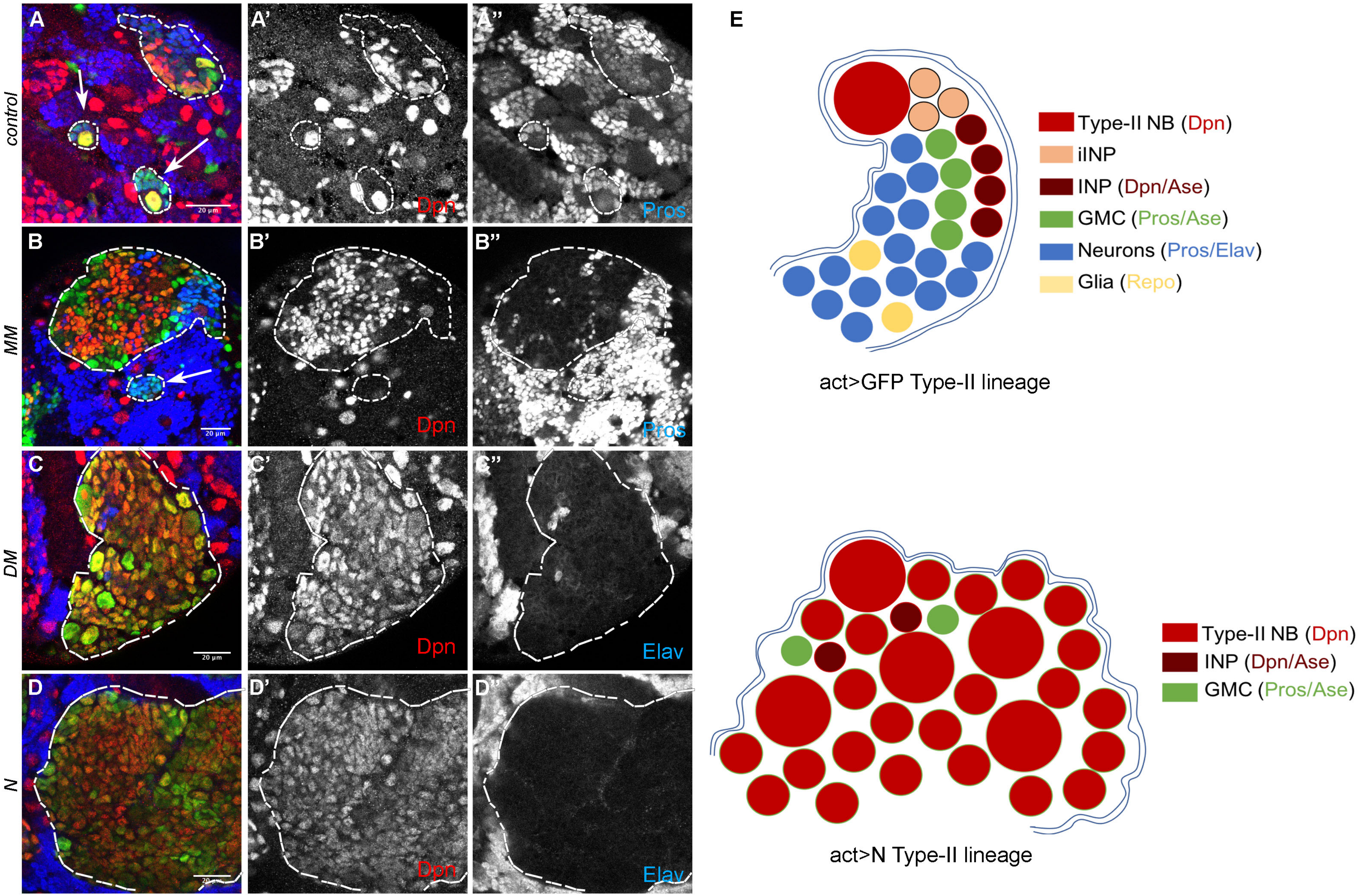

**Figure 1-Suppl2 Notch and Hes induced hyperplasia in Type II lineages.** Panels A-D are stained to reveal NSCs (Dpn, red) and GMCs/neurons (Pros, blue). FLP-out clones are marked with GFP (green) and overexpress (A) GFP alone (control); (B) UAS-m8, UAS-mγ (MM); (C) UAS-dpn, UAS-mγ (DM); (D) UAS-NΔecd (N). Clones are outlined. In panels A and B adjacent smaller Type I clones are also outlined and marked with an arrow. (E) Cartoons depict the cellular composition of wt vs *N* overexpressing TypeII lineages. *MM* and *DM* lineages would be the same, only they should additionally have a few blue cells (neurons). Scalebars 20μm.

### Figure 1 - Supplement 3

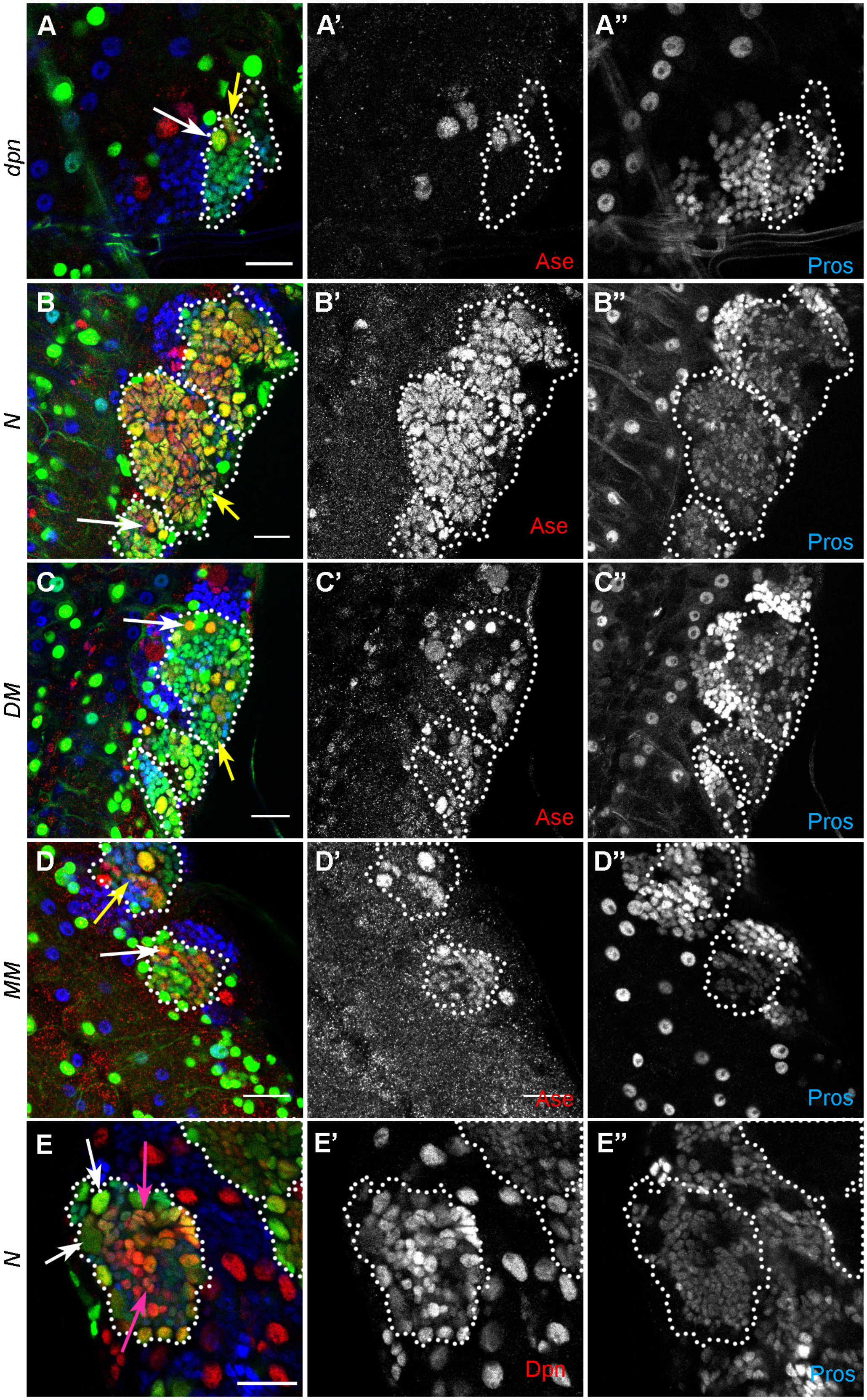

**Figure 1-Suppl3 Further examples of neural hyperplasia caused by N/Hes overactivity.** All panels show *act>STOP>Gal4* clones expressing GFP and transgenes as marked. A-D: NSCs/GMCs are detected by Ase (red); GMCs/young neurons are detected by Pros (blue) – individual channels are shown in greyscale. A is essentially wt, as *dpn* expression alone rarely causes weak defects in Type I lineages. White arrows mark examples of Ase positive/ Pros negative (NSC-like) cells. Yellow arrows mark double-positive (GMC-like) cells. E: Dpn (red) marks NSC-like cells (examples shown by white arrows) and Pros (blue) marks GMC/neuron-like cells. Examples of doubly Dpn/Pros positive cells are shown by magenta arrows. Scalebars 20µm.

### Figure 1 - Supplement 4

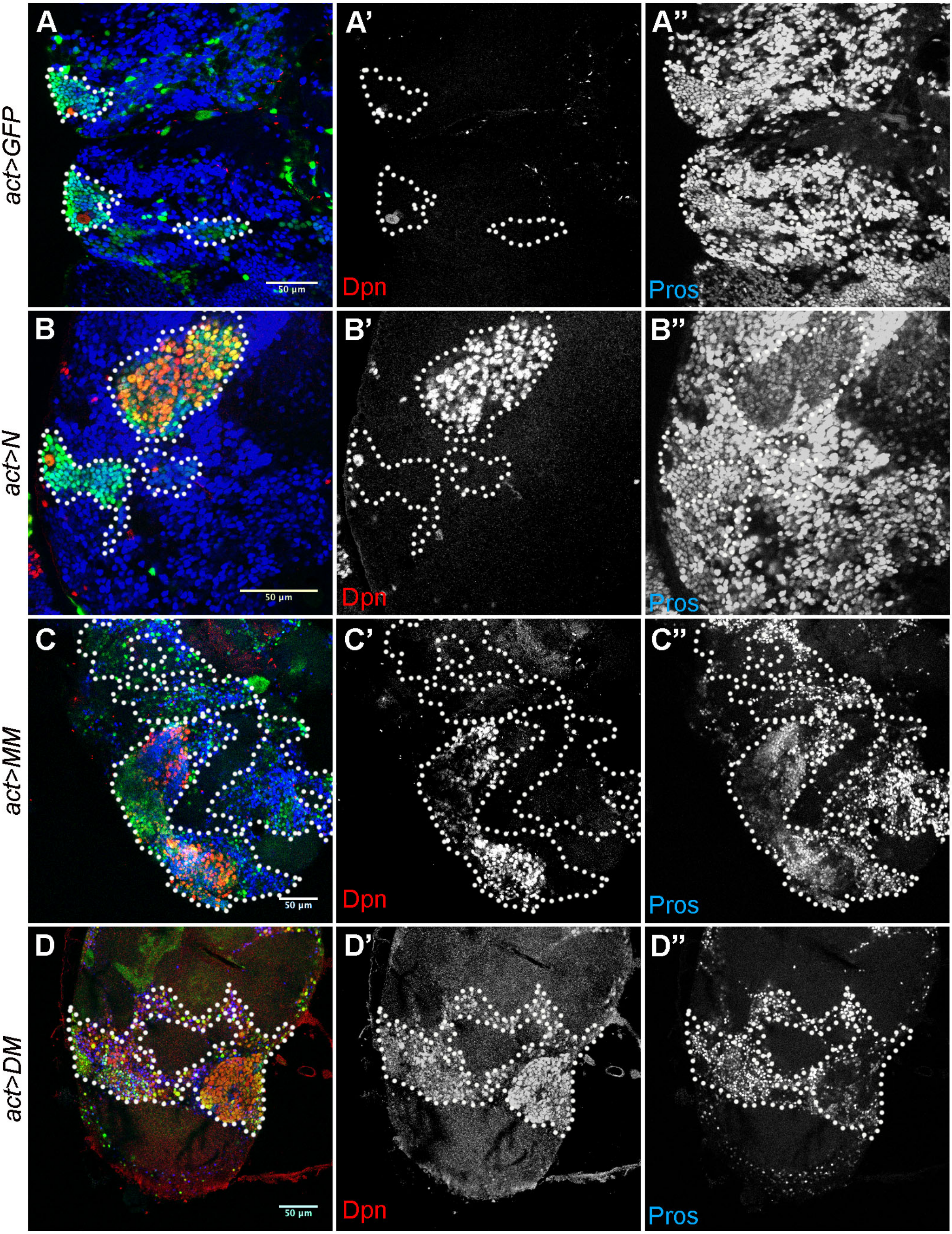

**Figure 1-suppl4 N/Hes NSC hyperplasias persist after pupariation.** A,B: brain lobes dissected at 24h APF; C,D: brain lobes dissected from freshly eclosed adult escapers. All animals expressed GFP together with the indicated transgenes from an act>STOP>Gal4 driver after hsFLP induction at early larval stages. Dpn is stained red and Pros blue. Individual channels shown in greyscale. The two clones that retain a Dpn-positive NSC in the wt (A) are mushroom body lineages that continue proliferating into the early pupal stages. Scalebars 50µm.

### Figure 3 - Supplement 1

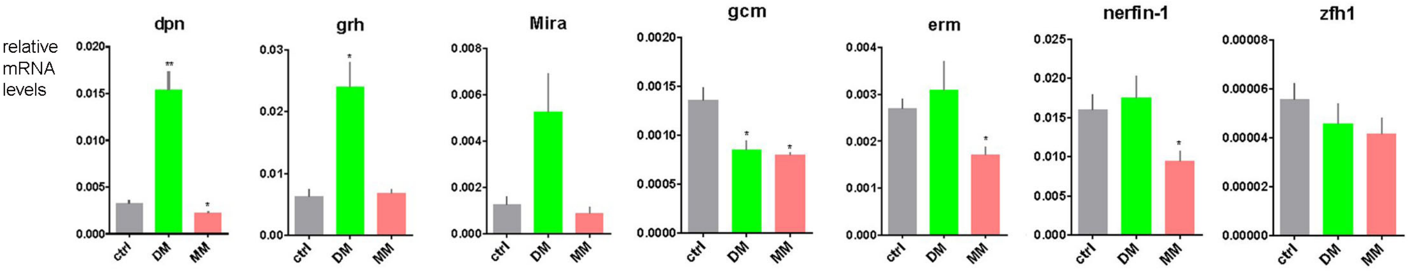

**Figure 3-Suppl1 qPCR validation of select mRNAs.** Expression levels were calculated by q-RT-PCR from *DM* or *MM* overexpressing CNSs (using *grhNB-Gal4*) or wt controls. Expression levels are shown relative to *RpL32* RNA. Error bars show the standard error of the mean from triplicate measurements. Asterisks indicate samples that are significantly different from control by Student's t-test (* P<0.05, **P<0.01).

#### Figure 3 – Supplement 2

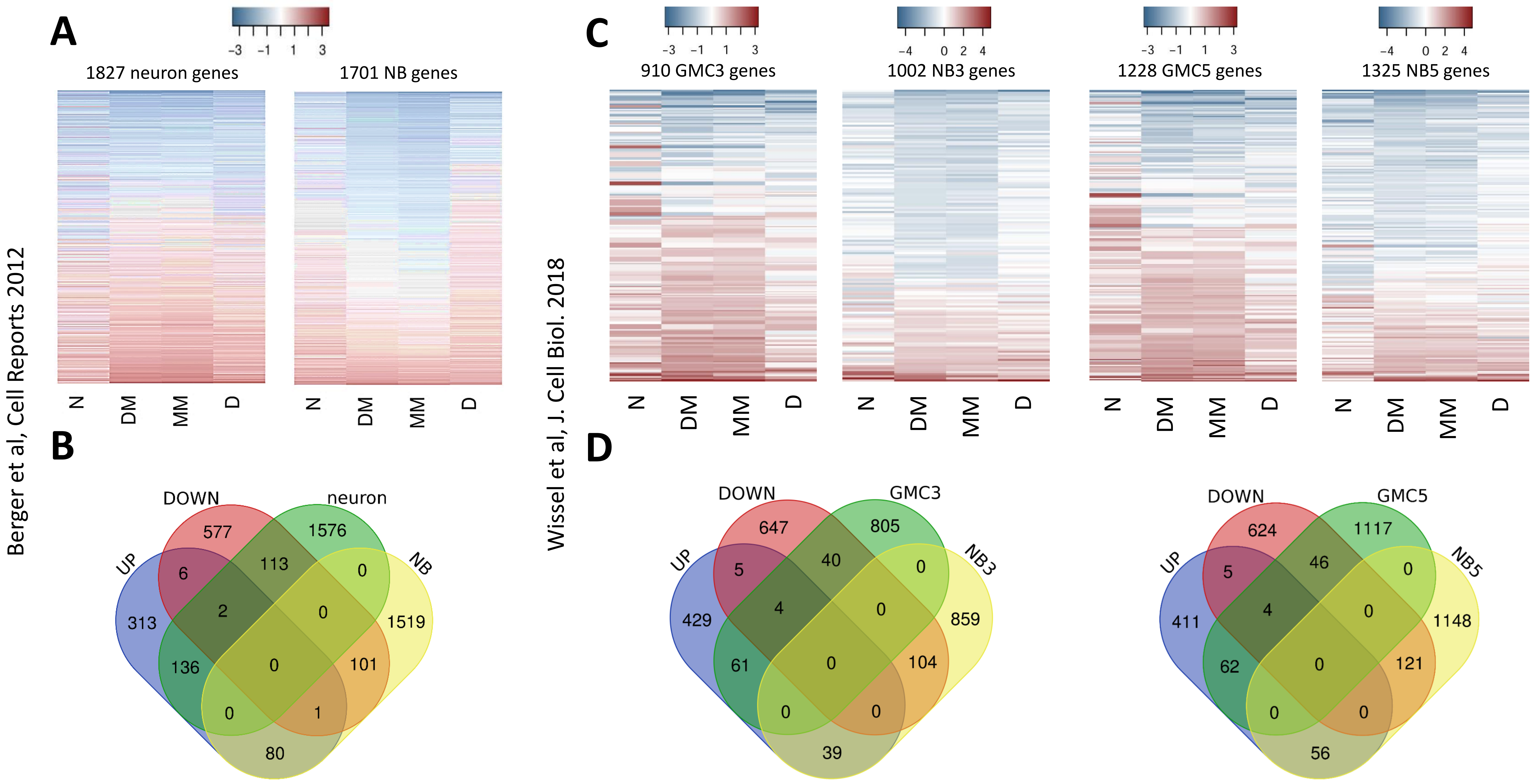

**Figure 3-Suppl2 Comparison of N/Hes differentially regulated genes with neuron, GMC and NSC-enriched gene-sets. (A,B)** Comparison with the neurons vs NBs enriched genes defined by Berger et al (2012): they show differential enrichment in FACS-sorted neuron or NSC populations from wt CNSs. (A) Heatmaps showing the $log_2$(fold-change) of neuron-enriched genes (left) and NSC-enriched genes (right) in the N and Hes overexpression backgrounds. (B) Venn diagram showing overlap of high confidence (FDR≤ 0.05) UP or DOWN-regulated genes in the N/Hes conditions (union of N with DM and MM, as shown in Fig. 3B,C) with the neuron vs NSC enriched gene-sets. (C, D) Comparison with the NSC vs GMC enriched genes defined by Wissel et al (2018): dissociated NSCs were cultured in vitro and their smaller GMC progeny FACS-sorted from the larger NSCs. 3 and 5 refer to the hours of culturing. (C) Heatmaps showing the $log_2$(fold-change) of GMC-enriched genes and NSC-enriched genes from each culture timepoint in the N and Hes overexpression backgrounds. (D) Venn diagram showing overlap of high confidence (FDR≤ 0.05) UP or DOWN-regulated genes in the N/Hes conditions (as in B) with the GMC vs NSC enriched gene-sets from the two culture timepoints.

**Figure 3 - Supplement 3**

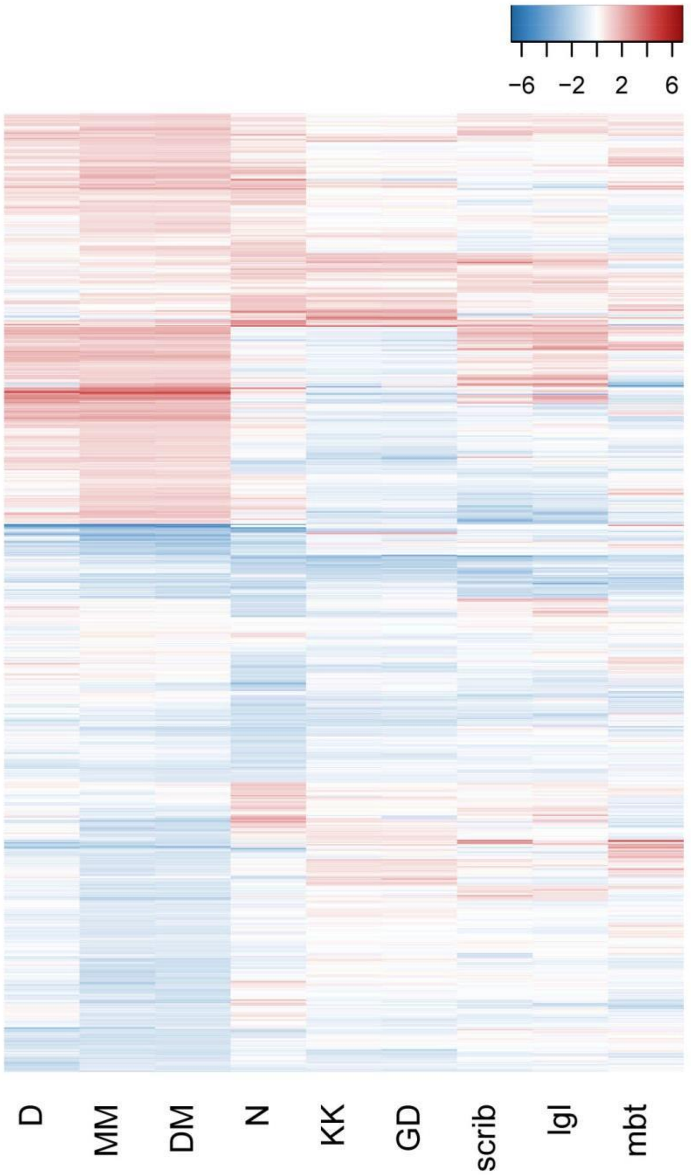

**Figure 3-Suppl3 Comparison of N and Hes CNS tumours with other CNS tumours. A.** Heat map of the fold-change of the 1410 genes selected by differential expression in the N or Hes tumours (Fig. 3A) in five additional tumour transcriptomes. KK and GD are two *brat* RNAi genotypes (Neumueller et al Cell Stem Cell 2011), whereas scrib, lgl and mbt refer to homozygous mutant larval CNSs for the genes *scrib*, *l(2)gl* and *l(3)mbt*, respectively, all of which show CNS hyperplasia.

#### Figure 5 - Supplement 1

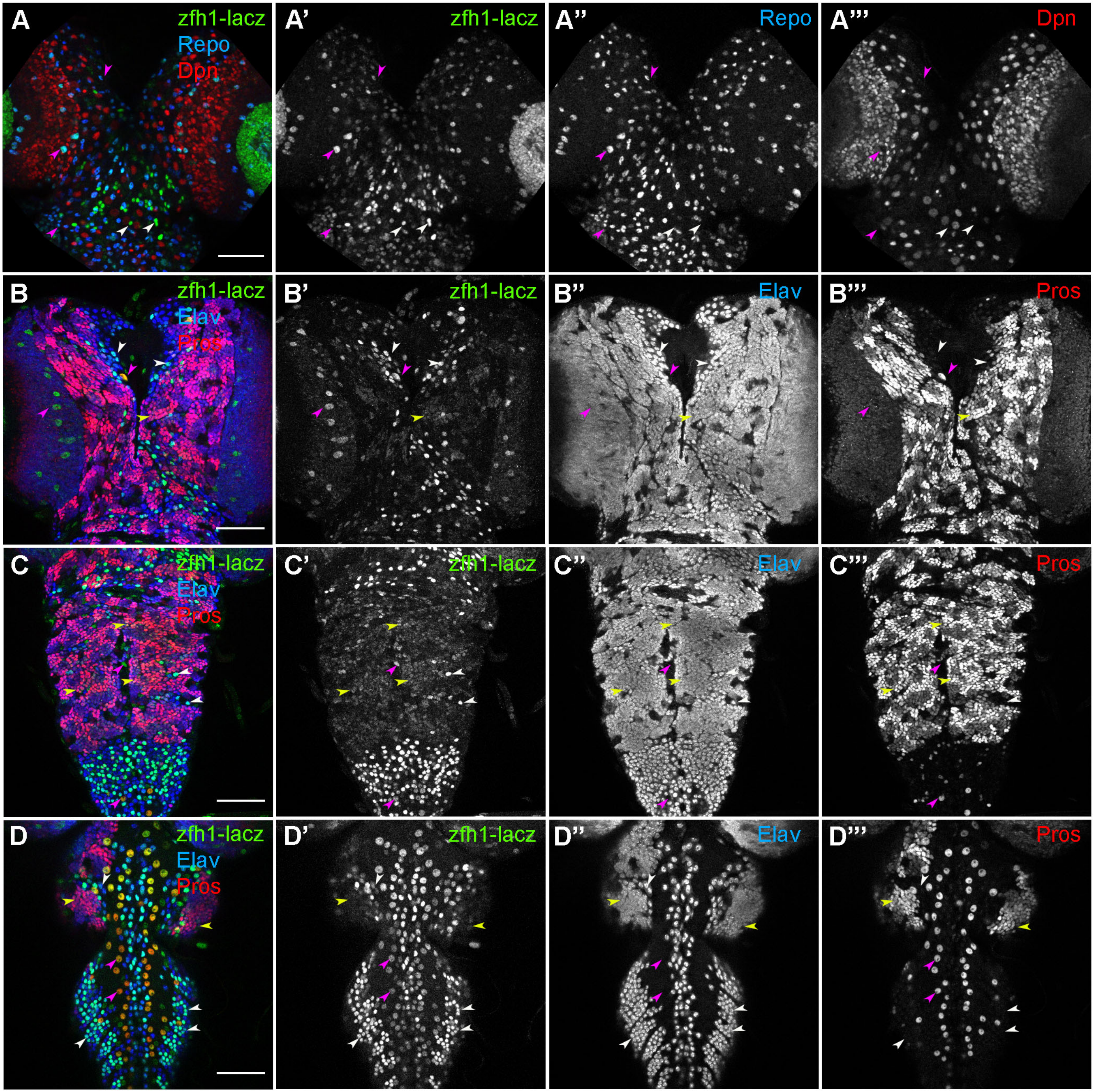

**Figure 5-Suppl1 *zfh1-lacZ* expression in the wt.** β-galactosidase is green in all panels. (A) Brain lobes counterstained with glial marker Repo (blue) and NSC marker Dpn (red). (B-D) Counterstaining for neuronal marker Elav (blue) and GMC/young neuron marker Pros (red) – note that Pros is also expressed in some glial cells. (B) Brain lobes. (C-D) VNC, ventral side (C) and dorsal side (D). Examples of GMCs/early neurons (Pros and Elav positive) are shown with yellow arrowheads, mature neurons (Elav positive/ Pros negative) by white arrowheads and glia (Repo positive) by magenta arrowheads. Scalebars 50µm

### Figure 6 - Supplement 1

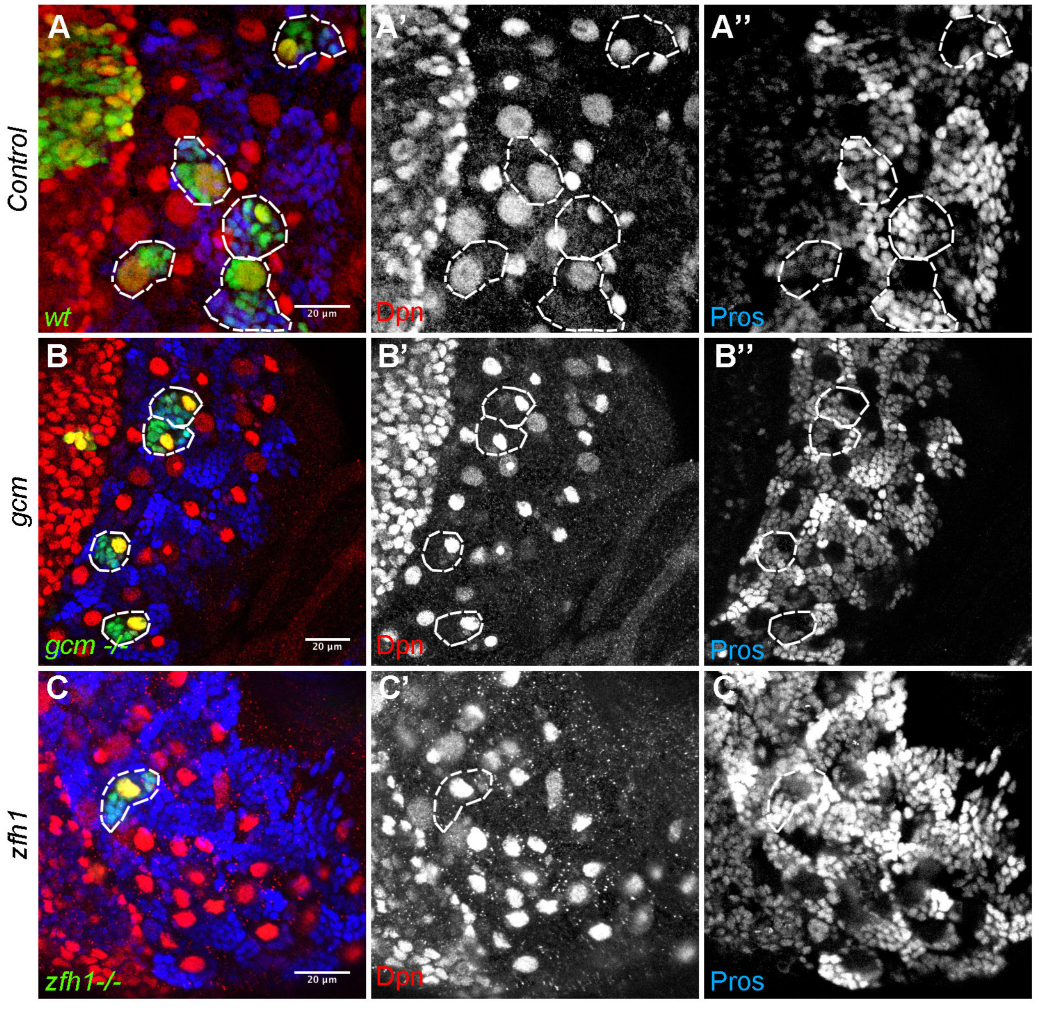

**Figure 6-Suppl1 Loss-of-function analysis of *zfh1* and *gcm***. MARCM clones are marked with GFP (green) and traced by dashed outlines. (a) WT, (B) clones are homozygous for *gcm*$^{N7\text{-}4}$ (C) clones are homozygous for *zfh1*$^{75.26}$. NSCs are imaged by Dpn (red) and GMCs/young neurons by nuclear Pros (blue). Individual channels shown in greyscale. Scalebars 20µm.

### Figure 7 - Supplement 1

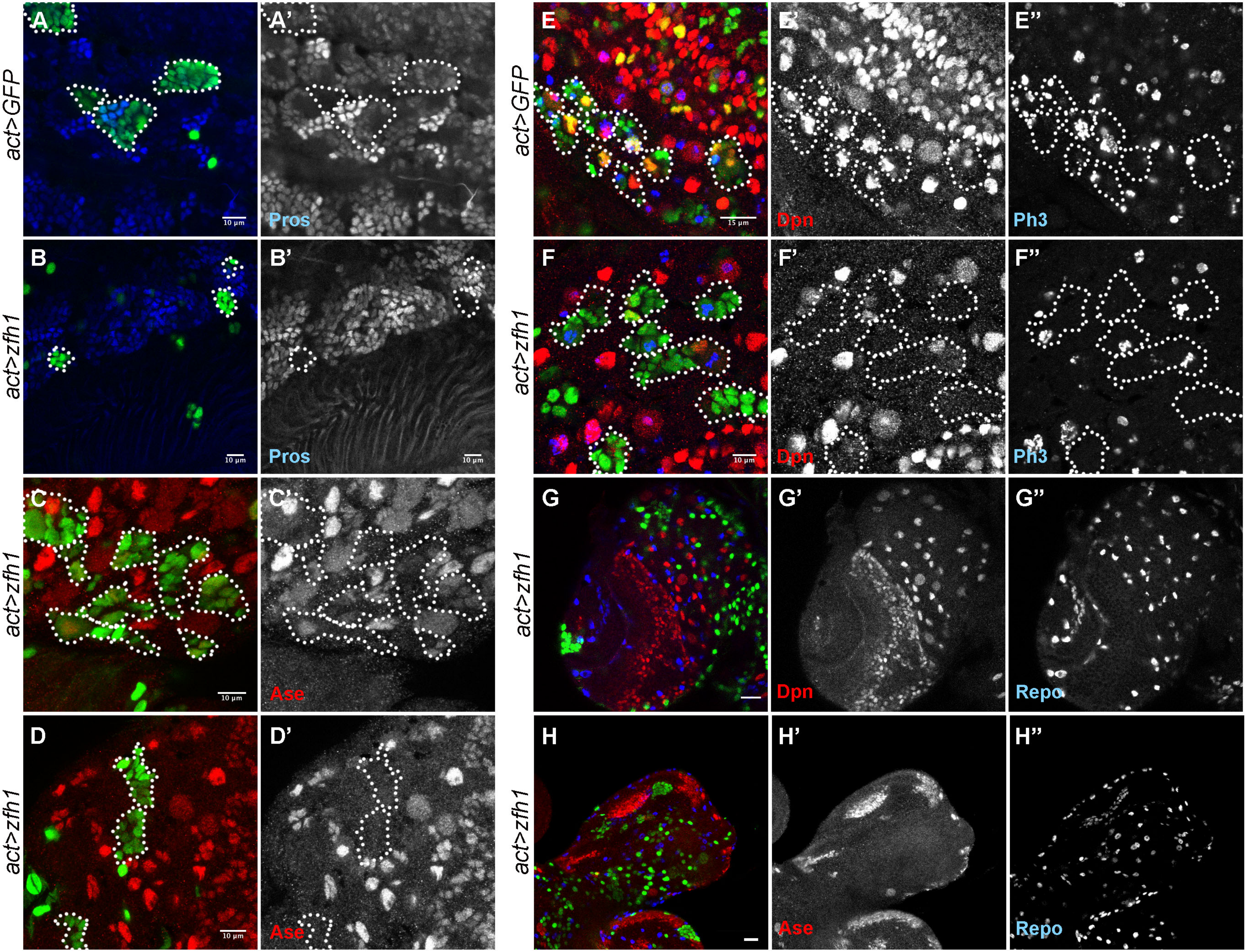

**Figure 7-Suppl1 Effect of *zfh1* misexpression on CNS lineages**. *act>STOP>Gal4* FLPout clones are marked by GFP (green) and traced with dotted outlines. (A) wt clones 3d ACI stained for Pros (blue, shown separately in A'). (B) *UAS-zfh1* clones 2d ACI stained for Pros (blue, shown separately in B'). (C,D) *UAS-zfh1* clones 1d (C) or 2d (D) ACI stained for Ase (red) shown separately in C', D'. (E) wt clones 1d ACI stained for Dpn (red) and the mitotic marker phosphor-H3 (blue). (F) *UAS-zfh1* clones 1d ACI stained as in E. (G,H) Examples of brain lobes carrying UAS-zfh1 clones 3d ACI and stained for the glia marker Repo (blue) and Dpn (red) in G or Ase (red) in H. G is a more superficial section than H. A-D scalebars 10µm, E scalebar 15µm, F scalebar 10µm, G-H scalebars 15µm.

### Figure 7 - suppl 2

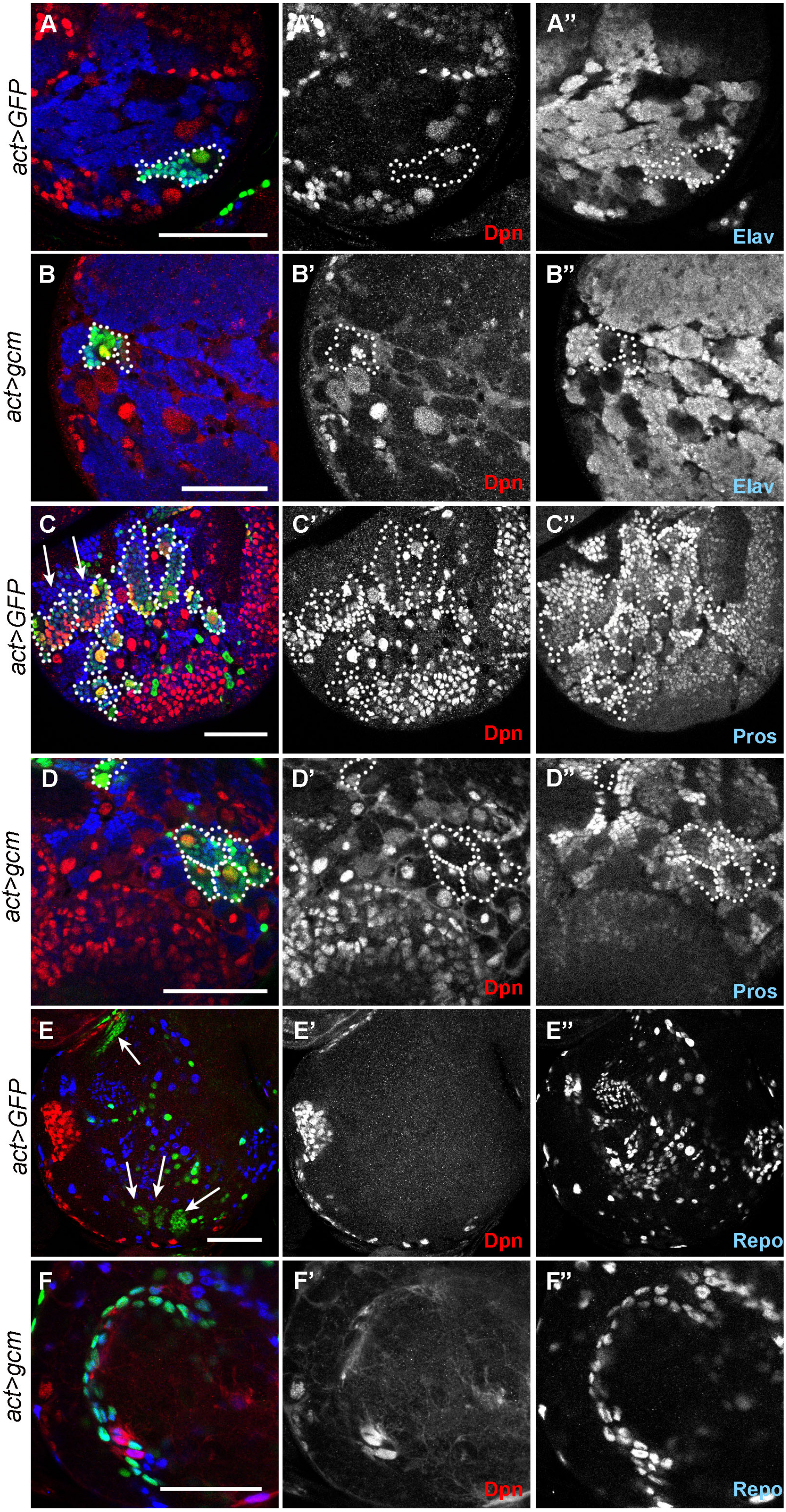

**Figure 7-suppl2 Effect of *gcm* misexpression on CNS lineages.** *act>STOP>Gal4* FLPout clones are marked by GFP (green) and traced with dotted outlines. (A,C,E) Control clones (GFP only); (B,D,F) UAS-gcm clones. (A,B) Sample stained for Dpn (NSCs, red) and Elav (neurons, blue). (C,D) Sample stained for Dpn (NSCs, red) and Pros (GMCs/ young neurons, blue); note two Type II clones in C (arrows). (E,F) Sample stained for Dpn (NSCs, red) and Repo (glia, blue). In E and F deep sections are shown where clone cells may have migrated away from the superficial stem cell (superficial lineages marked by arrows in E; note that the upper one derives from the OL); nevertheless in E there is hardly any GFP marking of glia, whereas in F most clonal cells have adopted a glial fate. All scalebars 50µm.

**Figure 8 - Supplement 1**

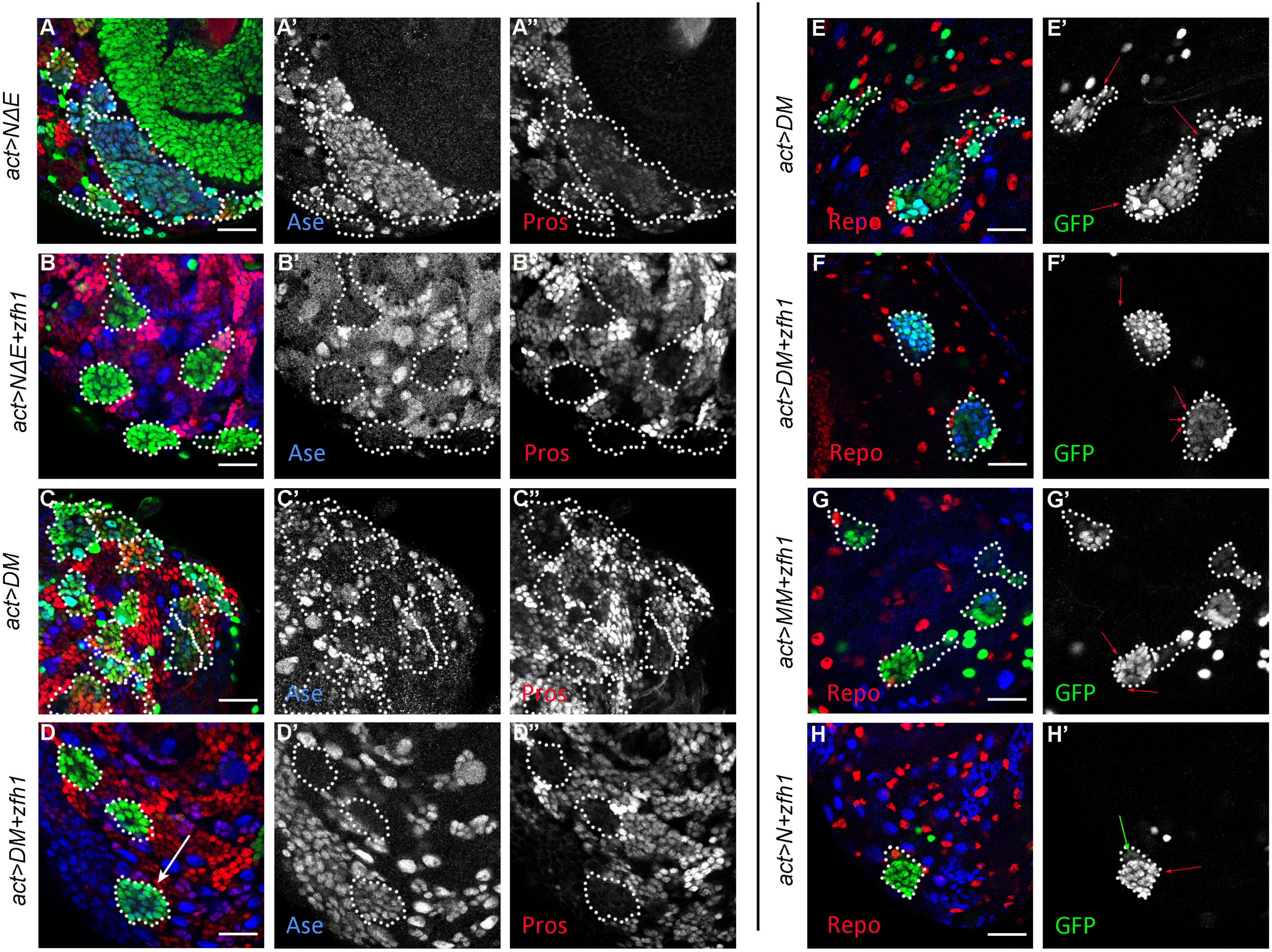

**Figure 8-suppl1 Effect of *zfh1* misexpression on *N*, *DM* and *MM* lineages**. *act>STOP>Gal4* FLPout clones are marked by GFP (green) and traced with dotted outlines. (A-D) Effects on Ase (blue) and Pros (red). Most Type I N/Hes clones lose Ase upon zfh1 expression, with some exceptions (arrow in D). (E-H) Effects on Repo. Even though there are several Repo positive cells adjacent to clones (red arrows), there are very few within clones (green arrow in H'). In E-H blue is Dpn. Scalebars 25µm.

### Figure 9 - Supplement 1

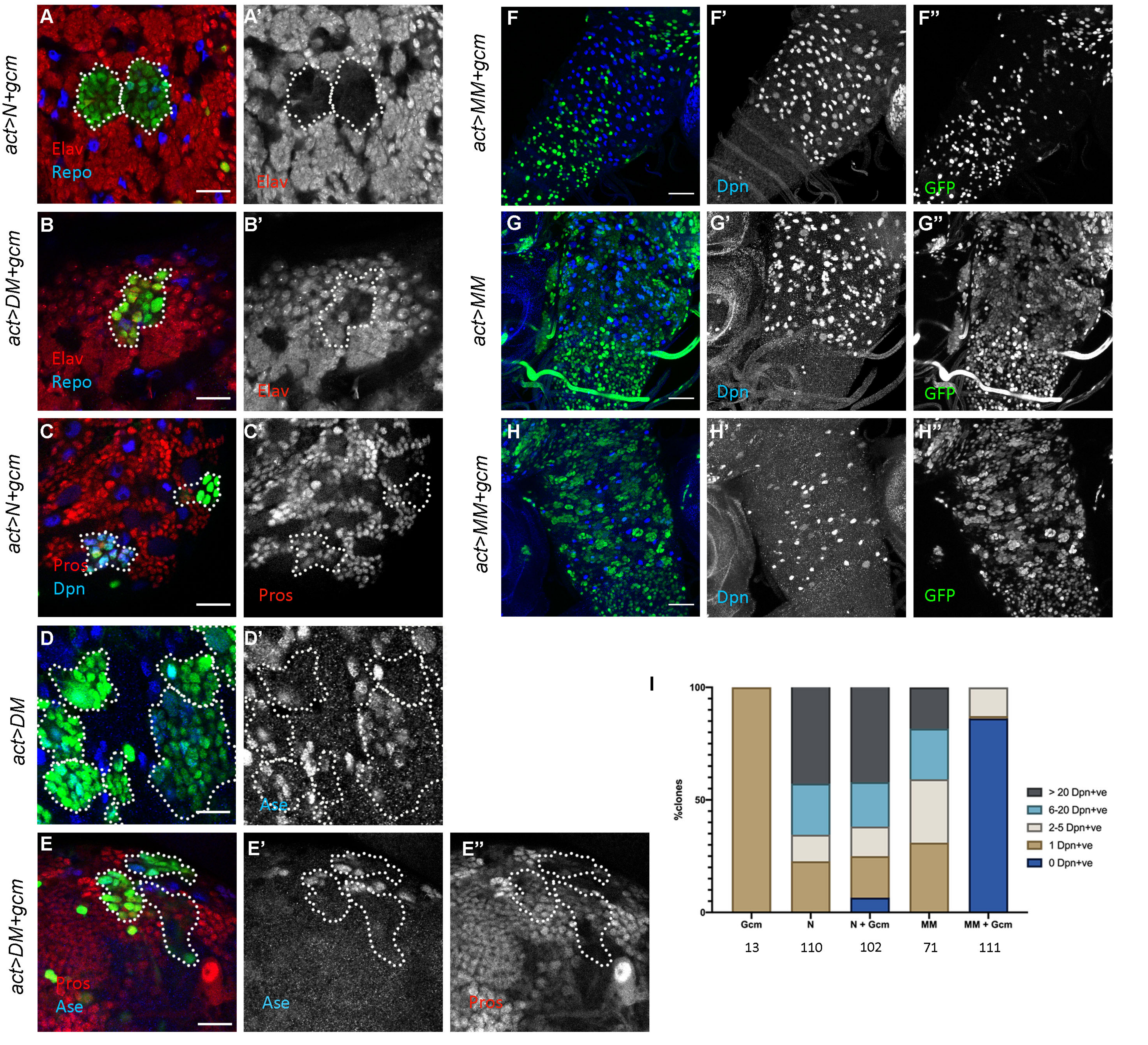

**Figure 9-suppl1 Effect of *gcm* misexpression on *N*, *DM* and *MM* lineages.** (A-E) *act>STOP>Gal4* FLPout clones are marked by GFP (green) and traced with dotted outlines. (A,C) misexpression of *NΔecd+gcm*. (B,E) misexpression of *DM+gcm*, (D) misexpression of *DM* (control for E). (F-H) Effect of *gcm* on the induction of Dpn (blue) by MM. F has no lineage clones (so the NSC number is not affected), whereas G and H have a large number of clones (see GFP-only channel in F"-H"). (I) Clones of the indicated genotypes were scored for the number of NSCs per clone. The total number of Type I clones scored for each genotype is shown below the chart. Scalebars: A-E 25µm, F-H 50µm.
